# Human Aging DNA Methylation Signatures are Conserved but Accelerated in Cultured Fibroblasts

**DOI:** 10.1101/605295

**Authors:** Gabriel Sturm, Andres Cardenas, Marie-Abèle Bind, Steve Horvath, Shuang Wang, Yunzhang Wang, Sara Hägg, Michio Hirano, Martin Picard

## Abstract

Aging is associated with progressive and site-specific changes in DNA methylation (DNAm). These global changes are captured by DNAm clocks that accurately predict chronological age in humans but relatively little is known about how clocks perform *in vitro*. Here we culture primary human fibroblasts across the cellular lifespan (∼6 months) and use four different DNAm clocks to show that age-related DNAm signatures are conserved and accelerated *in vitro*. The Skin & Blood clock shows the best linear correlation with chronological time (r=0.90), including during replicative senescence. Although similar in nature, the rate of epigenetic aging is approximately 62x times faster in cultured cells than in the human body. Consistent with *in vivo* data, cells aged under hyperglycemic conditions exhibit an approximately three years elevation in baseline DNAm age. Moreover, candidate gene-based analyses further corroborate the conserved but accelerated biological aging process in cultured fibroblasts. Fibroblasts mirror the established DNAm topology of the age-related *ELOVL2* gene in human blood and the rapid hypermethylation of its promoter cg16867657, which correlates with a linear decrease in ELOVL2 mRNA levels across the lifespan. Using generalized additive modeling on twelve timepoints across the lifespan, we also show how single CpGs exhibit loci-specific, linear and nonlinear trajectories that reach rates up to −47% (hypomethylation) to +23% (hypermethylation) per month. Together, these high temporal resolution global, gene-specific, and single CpG data highlight the conserved and accelerated nature of epigenetic aging in cultured fibroblasts, which may constitute a system to evaluate age-modifying interventions across the lifespan.

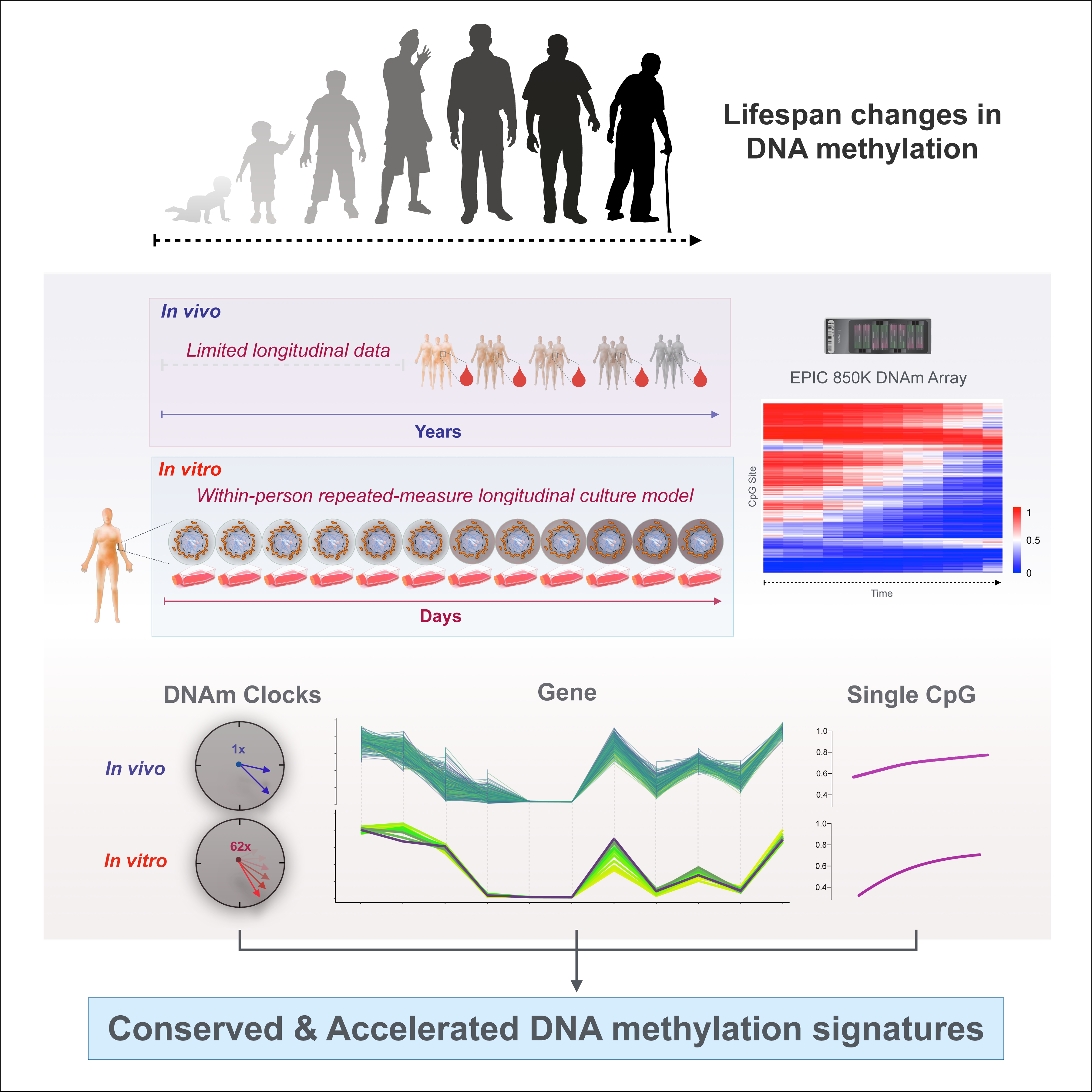
Graphical Abstract

## Introduction

As we attempt to develop increasingly precise and sensitive biological age indicators to detect and predict disease risk, a particular epigenetic marker that has gained interest is methylation of cytosine nucleotides in DNA (DNAm) (Horvath & Raj, 2018). DNAm is a stable epigenetic mark that serves to establish cell lineages during differentiation (Meissner et al., 2008) but rapid changes in DNAm are also known to occur on the time scale of days and weeks, including circadian oscillations in cells and animals (Oh et al., 2018), and over years with cellular aging (Wilson & Jones, 1983). In human tissues, specific cytosine-guanine dinucleotides (CpGs) undergo stereotypic loss (hypomethylation) or gain (hypermethylation) of methylation with advancing chronological age. Notably, among the CpGs most consistently found to undergo hypermethylation with age is *ELOVL2* cg16867657 (Garagnani et al., 2012; Gopalan et al., 2017; Johansson, Enroth, & Gyllensten, 2013; Wang et al., 2018), indicating the presence of site-specific alterations in DNAm with age in human populations. However, despite the availability of longitudinal DNAm datasets in human blood, it has remained challenging to track dynamic changes in DNAm across the entire human lifespan and to evaluate the effects of age-modifying genes or environmental perturbations without appropriate laboratory-based experimental systems.

In human tissues, epigenetic biomarkers, or biological “clocks” have been developed that reliably track chronological age (Horvath & Raj, 2018). DNAm clocks use penalized linear regression models, such as elastic-net regression to select CpGs whose change in DNAm correlate (either positively or negatively) with chronological age. For example, the Horvath pan-tissue clock (Horvath, 2013) and the Hannum clock (Hannum et al., 2013) have been widely used in epidemiological studies. They provide moderate to excellent estimates of chronological age (r>0.90) in human tissues (Horvath & Raj, 2018). Subsequent minimalist algorithms have also shown that the methylation levels of only three to eleven CpGs from human blood can provide fairly accurate age predictions (mean absolute error 2.72 - 5.4 years) or predict mortality (Li, Li, & Xu, 2018; Weidner et al., 2014; Zhang et al., 2017). However, little is known about the kinetics of DNAmAge and DNAm levels at single-CpG resolution across the human lifespan, in part because it is difficult to repeatedly obtain biological samples from a given individual across a sufficiently long period to robustly characterize DNAm kinetics.

Primary human fibroblasts are derived from minimally invasive skin biopsies, can be grown *in vitro*, and have been used as an experimental model of aging (Tigges et al., 2014). Fibroblasts divide at a regular rate and undergo a specific number of division (i.e., the Hayflick limit) before reaching senescence, a state characterized by replicative arrest and metabolic remodeling (van Deursen, 2014). In this system, senescence can also be induced by genotoxic stress, such as irradiation and DNA damage (Correia-Melo et al., 2016), providing an appealing model to study factors that influence the biological aging process. Previous studies have identified CpGs whose methylation levels distinguish cultured senescent from younger cells (Franzen et al., 2017; Koch et al., 2012), and work has provided preliminary evidence that DNA methylation age (DNAmAge) algorithms may be sensitive to replicative and radiation-induced senescence (Horvath et al., 2018; Kabacik, Horvath, Cohen, & Raj, 2018; Lowe, Horvath, & Raj, 2016). Although fibroblasts from older donors show reproducible differences in metabolic function and transcriptional state (Braam et al., 2006; B. D. Johnson, Page, Narayanan, & Pieters, 1986), it is unclear whether age-related DNAm changes in primary cultured cells reflect a distinct state unique to *in vitro* conditions, or if they replicate the ubiquitous molecular signatures that occur in aging human tissues.

Here, we examine to which extent different human DNAm clocks and age indicators track cellular aging in primary cultured fibroblasts, and compare the rate of aging *in vivo* and *in vitro*. We use repeated DNAm measurements across the cellular lifespan to define the spectrum and rate of both linear and nonlinear, hypo- and hypermethylation kinetics of individual CpGs. Overall, our findings reveal that DNAm aging signatures between primary cultured fibroblasts and human blood leukocytes are conserved and accelerated, suggesting that human fibroblast aging exhibits key molecular events that recapitulate human aging (Figure 1A). This study demonstrates that human biological aging in the order of decades can be mimicked with *in vitro* laboratory experiments over a few months.

**Figure 1.**
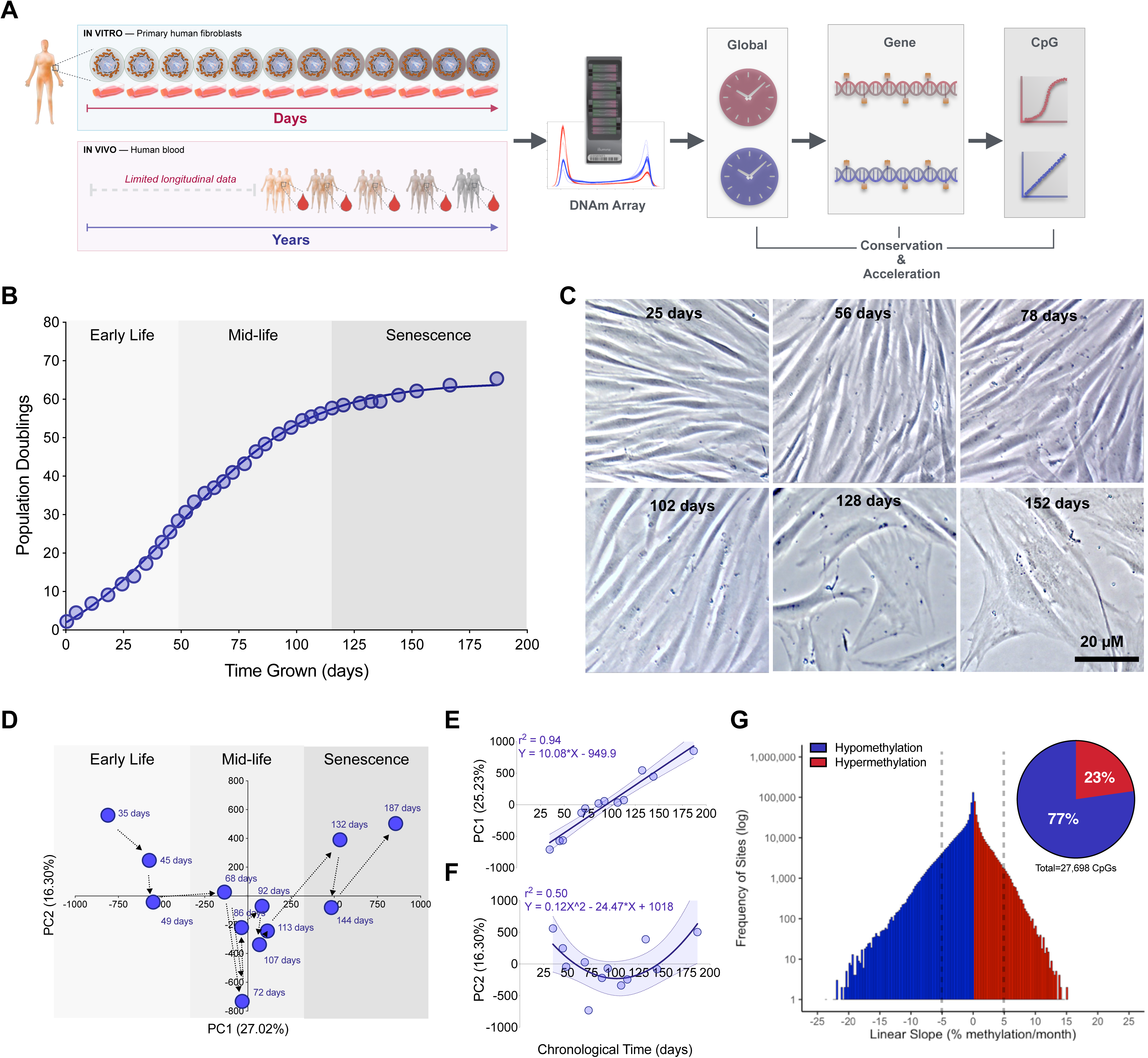
Primary human fibroblasts undergo global DNA methylation changes across the cellular lifespan. **(A)** Overview of study design, which includes 12 timepoints for a longitudinal assessment of DNAm across the cellular lifespan, and comparison with longitudinal data from human blood (Wang et al. 2018). **(B)** Cumulative population doublings for cultured primary dermal human fibroblasts. **(C)** Fibroblasts across different stages of the lifespan imaged by phase contrast microscopy after 5 days of growth at each respective passage. **(D)** Principal component analysis (PCA) of all CpG sites across the cellular lifespan; the first two components are plotted and each timepoint is indicated by the number of days in culture. **(E)** Correlation between the first and **(F)** second principal components and chronological age. **(G)** Frequency distribution of all CpG sites ordered by their rate of methylation change per month across the lifespan. Dotted lines indicate an arbitrary methylation rate threshold of 5%. Inset: Proportion of sites >5% per month that undergo age-related decrease (hypomethylation) or increase (hypermethylation) in DNAm.

## Results

### Primary human fibroblasts grown across the cellular lifespan

To examine the kinetics of DNA methylation across the lifespan, primary fibroblasts from a 29 year-old male were cultured and aged by continuous passaging until replicative senescence. The maximal linear rate of cell division was 0.75 divisions/day. In total, cells doubled approximately 56 times in 110 days, before reaching a plateau marking replicative senescence (Figure 1B). As expected, cellular senescence was associated with distinct cellular characteristics including loss of spindle-shaped cell morphology, appendage asymmetry, nuclear flatness, and reduced maximal confluency (Figure 1C).

### Whole methylome analysis across the cellular lifespan

DNA methylation was quantified for 866,736 CpG sites using the Illumina MethylationEPIC array at twelve timepoints collected approximately every 11 days across the cellular lifespan. When considering global DNAm variation for all CpGs together in a principal component analysis (PCA) (Figure 1D), the methylome appeared to follow a biphasic trajectory from early life, mid-life, and senescence. Combined, the first two principal components explained 41.5% of the variance in the methylome. Whereas the first component can be described as a linear function of chronological time (i.e., the amount of time cells were aged in culture) (r^2^=0.94, p<0.0001) (Figure 1E), the second component describes a quadratic function (r^2^=0.46, p = 0.068) (Figure 1F). These data strongly suggest large-scale, time-dependent linear and non-linear changes in the methylome with cellular age.

We then leveraged the high-temporal density of DNAm measurements across the lifespan and computed the overall rate (slope) of change in % DNAm per month, for each CpG (Figure 1G). Of the sites undergoing greater than 5% DNAm-change per month, 23% underwent hypermethylation, while 77% exhibited hypomethylation - yielding a skewed distribution towards hypomethylation. Consistent with previous work demonstrating age-related global hypomethylation (Unnikrishnan et al., 2018), a global analysis of the average DNAm values across all 867k CpGs also showed a linear decrease with age (r^2^=0.52, p = 0.0081), yielding a global 2.3% decrease in methylation (Figure S1C).

### DNA methylation clocks track age in cultured fibroblasts

To the extent that the age-related patterns of DNAm would be conserved between *in vivo* (i.e., human tissues) and *in vitro* (e.g., cultured human fibroblasts) (Horvath et al., 2018), we would expect algorithms trained on human tissues to also linearly capture the passage of time in cultured cells. On the other hand, if algorithms did not track cellular age *in vitro*, it would suggest that cell aging and human aging involve epigenetically distinct processes. To test if DNAm based aging clocks accurately track aging in cultured cells, we tested four independent clocks, all of which were trained and validated in cross-sectional human studies of various tissue types.

The four clocks are: (1) the Pan-Tissue clock developed as its name indicates on 51 different tissues types including cancer samples and consisting of 353 CpGs (Horvath et al. 2013); (2) the Skin & Blood clock trained on fibroblasts, keratinocytes, buccal cells, endothelial cells, lymphoblastoid cells, skin, blood, and saliva, and comprising 391 CpGs (Horvath et al., 2018); (3) the PhenoAge clock trained on whole blood and consisting of 513 CpGs (Levine et al., 2018); and (4) the Hannum clock also trained on whole blood and comprising 71 CpGs (Hannum et al., 2013). See methods for full clock descriptions. All clocks are strongly correlated with chronological age when tested in human tissues (r^2^=0.91-0.99).

In our lifespan fibroblast model, the Pan-Tissue Clock captured a linear increase in DNAmAge of 14 years in 78 days of growth (r^2^=0.83), but failed to capture chronological aging past 110 days in late life where little to no cellular division occurs over a period of approximately two weeks (i.e., replicative senescence) (Figure 2A). In contrast, the Skin & Blood clock returned a linear increase of 25 biological years over the entire lifespan of 152 days (r^2^=0.90) (Figure 2B). Other clocks trained solely on whole blood, the PhenoAge and Hannum clocks, were also able to predict a linear increase in biological age in cultured fibroblasts (r^2^=0.85, 0.76 respectively), but also failed to capture an increase in DNAmAge during replicative senescence (Figure S2A-B). The Skin & Blood clock was the only clock able to capture aging linearly through replicative senescence, suggesting that the Skin & Blood clock may be a better age estimator for *in vitro* aging (Horvath et al., 2018).

**Figure 2.**
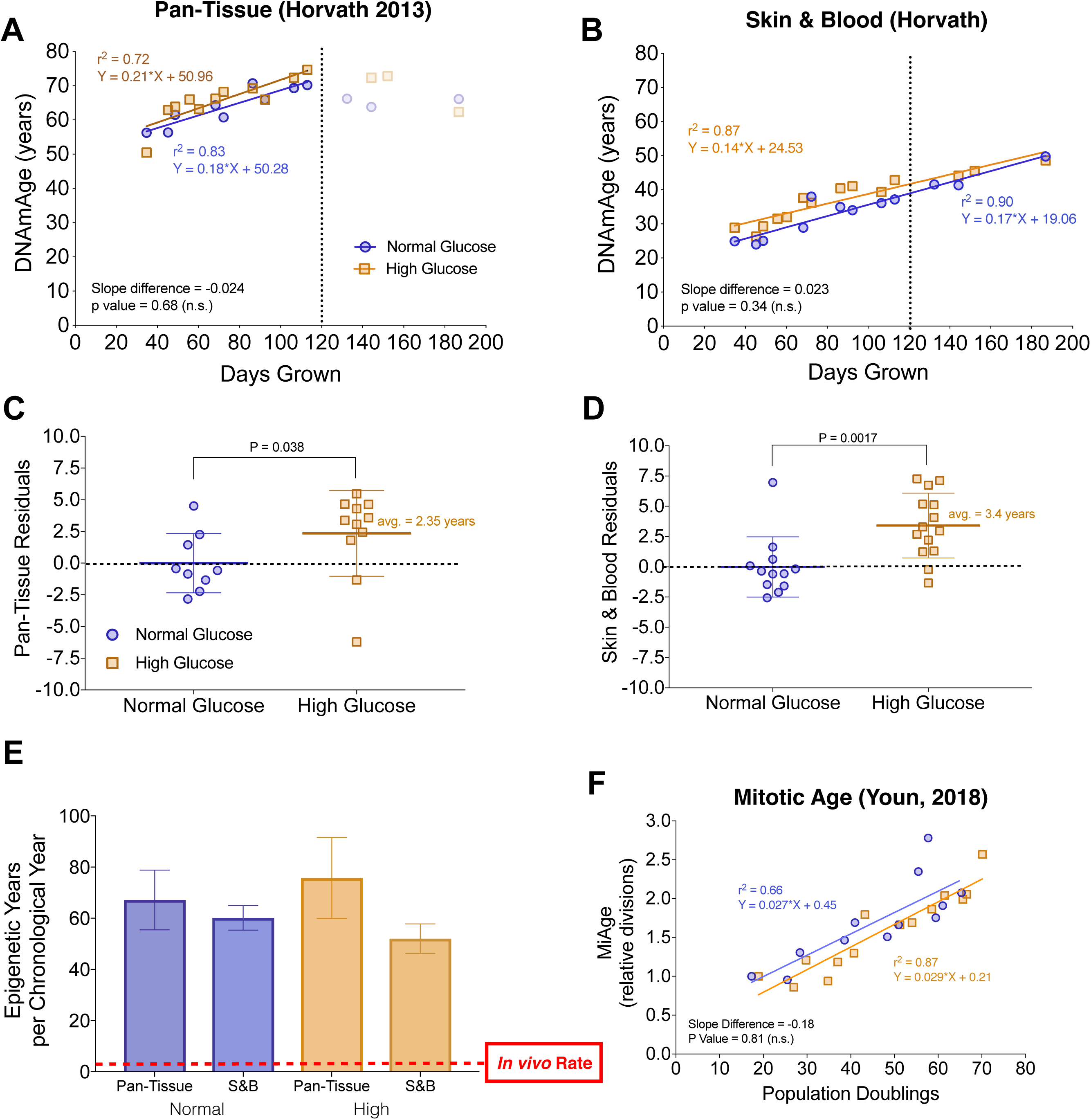
DNAm clocks track aging across the cellular lifespan, are sensitive to glucose levels, and reveal accelerated aging in vitro. **(A)** Linear regression of chronological age (e.g., days in culture) and predicted DNAmAge in primary human fibroblasts using the Pan tissue (Horvath et al., 2013) and **(B)** Skin & Blood (Horvath et al., 2018) clocks. Cells were cultured either in normal glucose (n = 12 timepoints, 5.5mM) and high “diabetic” levels of glucose (n = 14 timepoints, 25mM). The dotted line indicates the estimated point at which division rate substantially decreases (i.e., replicative senescence). Note that the Skin & Blood clock remains linear throughout the lifespan. **(C-D)** Box plots of the residuals from regressions in A and B, comparing normal and high glucose cells across lifespan. Each datapoint reflects the residual score for each timepoint assesses; non-parametric unpaired Mann-Whitney test. **(E)** The rate of epigenetic aging measured as the slope of DNAmAge and chronological time in years over the linear portion of the regressions in A and B. For the regression of the Pan Tissue Clock, only timepoints in early- and middle- life are used. **(F)** Regression of MiAge estimated cell divisions to actual population doublings calculated from cell counts performed at each passage.

These different clocks have minimal overlap in their underlying CpGs (0-6% similarity, Figure S2H) and each of the clocks’ CpGs change relatively little over the lifespan (Figure S2G). Nevertheless, the consistent linear correlation between DNAmAge based on all four clocks and chronological age during early- and middle-age (r^2^=0.76-0.90) suggests that a human epigenetic aging signature is conserved between cultured cells and *in vivo* aging.

### Accelerated cellular aging in cultured human fibroblasts

To test whether *in vitro* fibroblast DNAmAge is sensitive to metabolic perturbations as reported in previous studies (Horvath et al., 2014), cells of the same individual were grown in parallel in either normal or hyperglycemic (i.e., diabetic, 25mM) glucose levels and DNAm measured at the same timepoints. DNAmAge estimates from both the Pan-Tissue and Skin & Blood clocks (Figures 2C-D), but not other clocks (Figures S2D-E), showed that chronic hyperglycemic conditions caused a significant elevation in the predicted age by 2.4 and 3.4 years, respectively. However, the rate of aging, reflected in the slope of time and DNAmAge, was not significantly elevated in any of the clocks. This age-acceleration provides proof-of-concept evidence that DNAmAge measured in primary human fibroblasts, as for some tissues *in vivo*, is also sensitive to environmental stressors.

Because DNAmAge algorithms were originally trained to yield one year of biological aging for each year of chronological time, the slope of the regression between chronological time and DNAmAge reflects the actual rate of aging. Thus, the slope can be compared between *in vivo* (slope = 1) and *in vitro* conditions. Strikingly, compared to aging in the human body, regardless of the clock used, cells grown *in vitro* showed a 60-65x (average = 62-fold) acceleration in the rate of epigenetic aging (Figure 2E).

Quantifying the association between the number of cell divisions and change in DNAmAge allows to back-calculate the rate of divisions expected to occur in the human body. We determined the number of cell divisions undergone during the linear growth phase (e.g., 30 population doublings) and divided that number by the epigenetics years elapsed (10 DNAmAge years), yielding the number of divisions that occur for each year. Based on this approach, we can estimate that in the body, cells undergo 2.8-3.2 divisions/year (average = 3.0 divisions/year), although this estimate requires empirical validation. In comparison, proliferative cultured fibroblasts divide at a rate of 181.9 divisions/year, again highlighting the accelerated nature of DNAm aging *in vitro*.

Further leveraging the direct assessment of cell divisions in culture, we had the opportunity to validate the Mitotic Age (MiAge) calculator originally developed from tissues of different ages (Youn & Wang, 2018). The MiAge calculator estimates the number of cell divisions using the stochastic replication errors acquired at individual CpGs in the epigenetic inheritance process during cell divisions. Here we confirmed that the MiAge calculator directly correlates with the actual number of cell divisions in culture (r^2^=0.66, p value = 0.0014, Figure 2F). Given that the MiAge calculator was built solely on DNAm datasets derived from tissues that have aged while in the organism, this finding further indicates that changes in the epigenetic landscape that occur during *in vivo* and *in vitro* aging are similar in nature.

### Gene analysis of ELOVL2 age-dependent DNA methylation and expression

To test the conservation of age-related DNAm at the gene level we then compared DNAm of candidate genes across the lifespan in both cultured fibroblasts and in blood from a longitudinal twin cohort study. The human blood cohort included 1,011 samples collected from 385 Swedish twins, with DNAm on the 450k array assessed up to five timepoints over a 20-year period.

In particular, we focused our analysis on *ELOVL2*, a well-validated gene showing age-related hypermethylation in epigenome-wide association studies for cg1686757 (Garagnani et al., 2012; Gopalan et al., 2017; Johansson et al., 2013; Kananen et al., 2016; Wang et al., 2018). cg1686757 is located ∽300bp from the transcription start site (TSS) in a CpG island within the *ELOVL2* gene promoter (Figure 3A). Interestingly, we found that the topology of DNAm levels at different CpGs along *ELOVL2* was nearly identical to that of fibroblasts (Figure 3B). Beyond these gene-wide similarities, the age-related changes in DNAm at single CpGs were also highly conserved in cultured fibroblasts. Specifically, cg1686757 was linearly hypermethylated by 21% *in vivo* (over 50 years), and correspondingly hypermethylated by 35% *in vitro* (over 98 days) followed by a plateau during replicative senescence (Figure 3C). To test whether changes in DNAm may regulate gene expression, we also performed RNA sequencing on cultured fibroblasts at three timepoints across early- and mid-life. The conserved age-related hypermethylation of *ELOVL2* cg1686757 was linearly correlated with a >80% reduction in ELOVL2 transcript levels with age (r^2^=0.87, Figure 3D). This combined transcriptomic and epigenomic data suggests that in this isolated fibroblast system where other confounding factors are avoided, hypermethylation of cg1686757 in promoter CpG island is associated with the expected gene repression.

**Figure 3.**
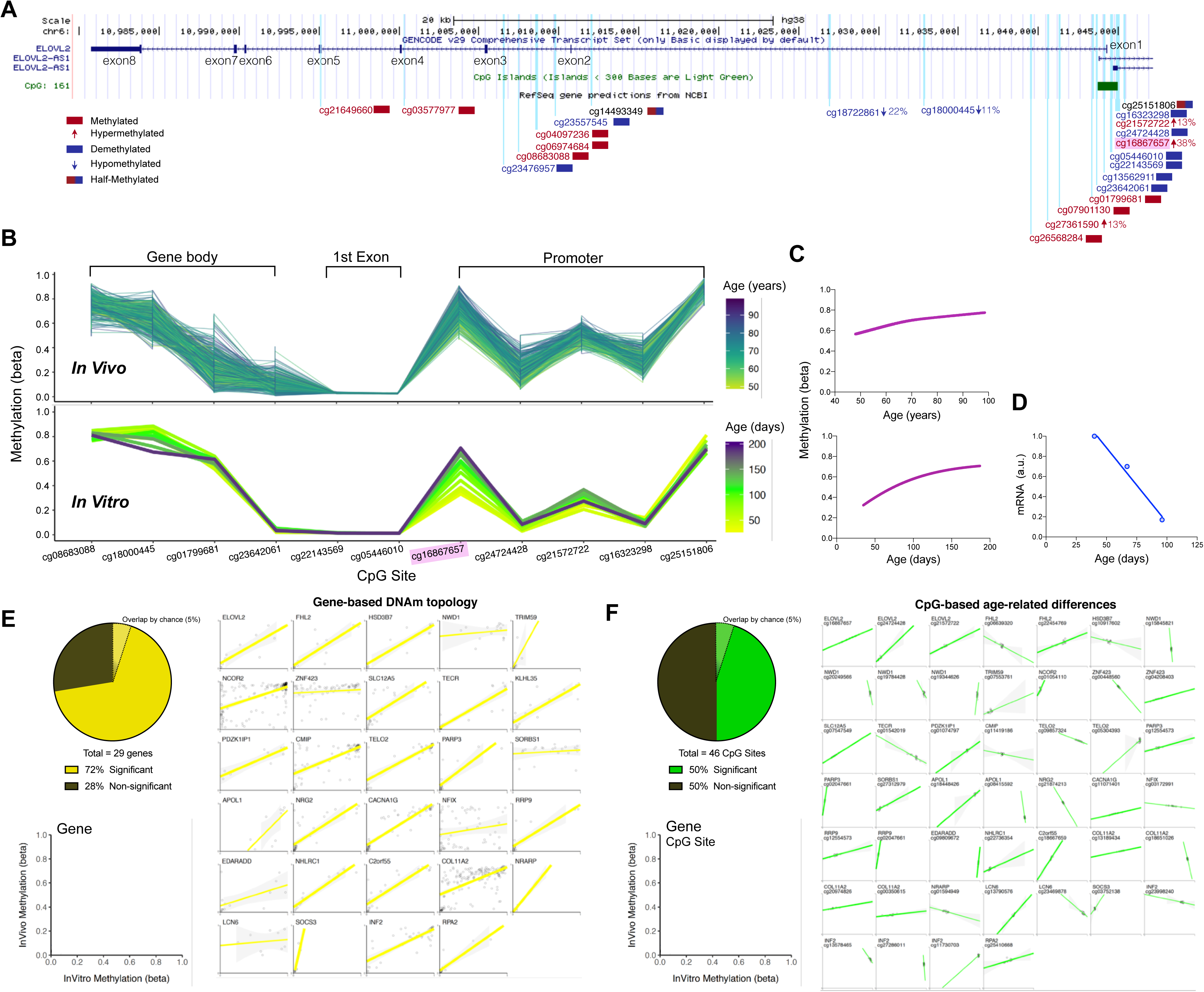
ELOVL2 DNAm topology and age-related hypermethylation and expression are conserved between human blood and cultured fibroblasts. (**A**) Overview of the *ELOVL2* gene with 8 exons, 7 introns, and 1 CpG Island (green box) located in the promoter region. All EPIC array CpG sites are mapped as vertical light-blue lines. CpGs with >75% (red) or <25% (blue) methylation levels are color coded, and CpGs exhibiting significant DNAm changes in human blood across the cellular lifespan are indicated by an arrow with their % change in DNAm level. (**B**) DNAm topology graph of *ELOVL2* CpG sites within the promoter, first exon, and gene body present in both the EPIC (*in vitro*, fibroblasts) and 450k (*in vivo*, blood) arrays. Methylation levels from whole blood (top,, Wang et al. 2018) and cultured human fibroblasts (bottom) are juxtaposed, highlighting their similar topology. In both datasets, each line represents a different individual (*in vivo*) or timepoint of the same individual’s cells (*in vitro*), color-coded by age. (**C**) Sigmoidal fit line of the DNAm levels for cg1686657 *in vivo* (top) and *in vitro* (bottom) across the lifespan.. (**D**) ELOVL2 transcript levels quantified by RNA sequencing across the early- and mid-life portions of the cellular lifespan (linear regression, n = 3 timepoints, r^2^ = 0.98, p value = 0.012). (**E**) Analysis of topological similarity, quantified as the correlation across all CpGs that map to a given gene between *in vivo* and *in vitro* systems. Each datapoint is the average methylation values across all ages/passages, plotted for each of the top 29 age-associated genes reported in Wang et al. (2018). Significant correlations (p < 0.05) are shown as thick regression lines, with 95% confidence interval (shaded area). Inset: proportion of *in vivo* - *in vitro* correlations that are significant correlations (p < 0.01). (**F**) Same as E but for 46 single CpGs whose methylation levels are positively correlated (p < 0.05) with age (Wang et al. 2018). Each graph is for a single CpG and each datapoint (n=12) reflects time in culture (x axis) and corresponding human ages (y axis). Full size correlation graphs can be found in Supplementary Figure S4.

We then expanded this analysis to a broader set of genes including the top 29 significantly age-associated genes from the longitudinal Wang et al. 2018 blood dataset (Figure S6). Of these genes exhibiting significant age-related alterations in DNAm *in vivo*, compared to the 5% proportion of genes expected to replicate by chance, the pattern of methylation was replicated in 76% of genes *in vitro* (p < 0.05, Figure 3E). Similarly, 50% of the CpG sites whose DNAm levels significantly correlated with age in human blood showed a similar pattern of change in cultured fibroblasts (Figure 3F). These conserved gene-wide topological DNAm signatures and age-related changes at single CpGs suggest a strong conservation of epigenetic signatures between human tissues and the cultured fibroblast system.

To further evaluate the concordance between DNAm and gene expression with age across cell types and aging environments, we compared the overlap of significant age-related CpGs taken from three independent datasets (i) *in vitro* fibroblasts in this study (n = 64,686 CpGs), (ii) whole blood from Wang et al. 2018 (n = 1,316 CpGs), and (iii) Horvath et al. (2018) *in vivo* fibroblasts (n=28,744 CpGs). Thirty-five significant CpGs were shared amongst the three datasets (Figure S4A and Supplemental Table S1). Of these shared sites, 86% exhibited hypermethylation with age in cultured fibroblasts (Figure S4B), 87% were located in CpG islands (Figure S4C), and 49% were within 1500bp of the TSS (Figure S4D). Furthermore, of the 35 overlapping CpGs, seven were associated with significant age-related change in transcript level, six of which are located in CpG islands in proximity to the TSS. Accordingly, five of these genes were upregulated and two were downregulated with age, including *ELOVL2* (Figure S4E), consistent with the notion that CpGs in islands near the TSS may influence gene expression.

### Single CpG kinetics for age-related changes in DNA methylation

In cells aged *in vitro*, environmental conditions can be controlled while longitudinal repeated measures are serially collected. Thus, the effect of time (i.e., independent variable) can most directly be examined without the many confounding variables that naturally arise in humans. This system thus enable to map precisely DNAm changes of individual CpGs across the entire lifespan, which our gene-based analysis suggested were of substantial magnitude, non-linear across the lifespan, and possibly functionally significant.

To examine and visualize the diversity of individual CpG site kinetics across the cellular lifespan, we first used a generalized additive model (GAM) to estimate the degree of freedom that best characterizes the lifelong trajectory of each CpG on the EPIC array. We extracted the 1,000 sites with the lowest p values (all p < 6.0^-45^) (Figure 4A-B), classified them into degrees of freedom 1 (constant or linear change), 1-3 (quadratic), or 3-4 (cubic), and dichotomized sites by their direction of change with age, as either increasing (hypermethylation) or decreasing (hypomethylation).

**Figure 4.**
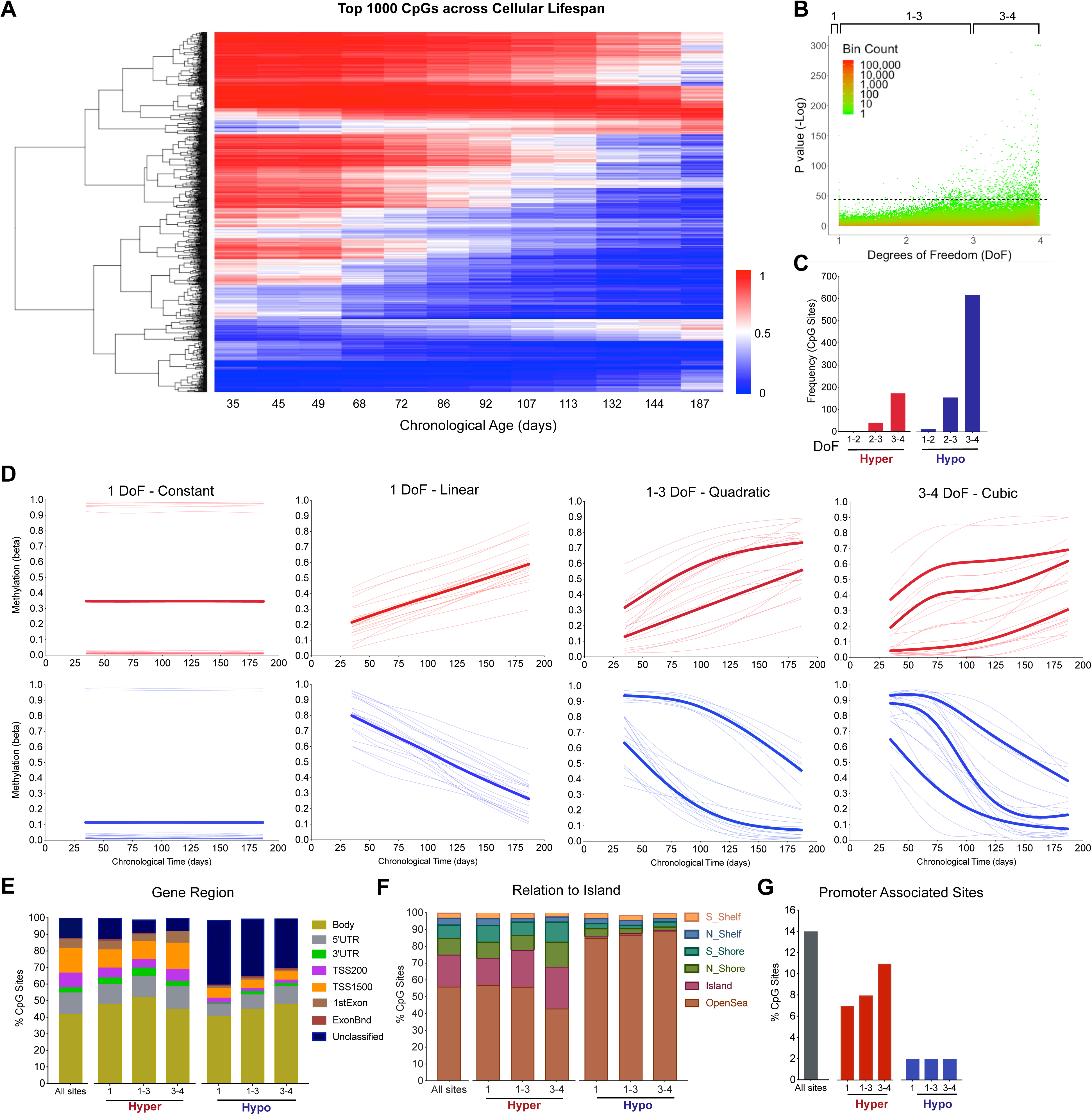
Lifespan trajectories of single CpGs reveal rapid linear and non-linear age-related changes in DNA methylation. **(A)** Heatmap of the top 1,000 age-related CpGs with the lowest P values from the generalized additive models (GAM) analysis across the lifespan (see methods for details). Hierarchical clustering using complete linkage and Euclidean distance [sqrt(sum((xi - yi)^2)))]. **(B)** P values for all EPIC CpG sites arranged by degree of freedom (DoF). Coloring scheme indicates datapoint density (log scale). **(C)** Proportion of the top 1,000 age-related CpGs that decrease (Hypo) or increase (Hyper) in DNAm levels with age, organized by DoF. **(D)** Example of lifespan trajectories (fitted models) using the top 20 most significant CpG sites undergoing hypermethylation (top) and hypomethylation (bottom), for DoF 1-4. Bolded lines represent the average of all sites with similar trajectories. **(E)** Distribution of the top 1,000 sites by gene regions **(F)**, relative to CpG islands, and **(G)** promoter region by DoF categories. The label “All sites” corresponds to all sites included in the EPIC array and is used as reference to evaluate the enrichment in specific genomic locations.

Consistent with our PCA results, this categorization revealed that more than 90% of the age-sensitive methylome varies non-linearly (degrees of freedom 2-4) across the lifespan (Figure 4C). Moreover, fitting non-linear GAM models to each CpG provided accurate estimates for both the magnitude and rate of age-related DNAm changes (Figure 4D). Sigmoidal hypomethylated sites showed magnitude of changes reaching up to 81% (e.g., from 98% in early life to 17% in senescence). The maximal rate of demethylation observed was as rapid as 47% per month (or more than 1.5% per day) compared to 15% per month for hypermethylated sites. Moreover, consistent with the fact that age-associated CpG sites from epigenome-wide association studies (EWAS) predominantly undergo hypomethylation (Day et al., 2013; Heyn et al., 2012), we found that the top 1,000 sites contained 2.7-fold more CpGs with hypomethylation relative to those undergoing hypermethylation. Thus, hypomethylation is both overrepresented across the methylome and occurs at a faster rate than hypermethylation during cellular aging.

Analysis of genomic location for sites undergoing hypo- and hypermethylation showed that although hypermethylated sites were not enriched in any genomic location, the prevalence of hypomethylated sites differed across loci. When stratified by DoF and compared to all CpGs surveyed on the EPIC array, hypomethylating CpGs were enriched by 1.5-2.0-fold in unclassified intergenic regions and by 0.5-0.6-fold in open seas. On the other hand, there was a 95% and 92% loss of enrichment for sites undergoing hypomethylation in CpG islands and gene promoters, respectively (Figure 4E-G), consistent with a relatively greater prevalence of hypermethylation in gene promoter CpGs - such as ELOVL2 cg1686757. This indicates the non-random distributions of age-related CpGs.

Interestingly, with the high temporal information in this system, the GAM model also identified some non-linear CpGs that showed multi-directional changes in methylation with age (Figures S5A-C). In these cases, a given CpG undergoes hypomethylation at one stage of the lifespan, followed by a change towards hypermethylation (i.e., biphasic), or vice versa. This finding demonstrates that although cellular aging can be successfully predicted on the basis of linear DNAm changes that constitute epigenetic clocks, this lifespan cellular system reveals the existence of nonlinear monotone and possibly bi-phasic DNAm trajectories at specific genomic locations.

## Discussion

There is a need to develop experimental models where cellular aging in humans can be studied across the lifespan. In this study, the global clock-based findings, gene-based profiling, and single CpG methylation kinetics among *in vitro* and *in vivo* conditions collectively highlight the conserved but accelerated epigenetic aging in cultured primary human fibroblasts. In just five months of cellular growth, the Skin & Blood clock tracked 25 years of biological aging. Some epigenetics clocks were also sensitive to metabolic stress, resulting in accelerated DNAmAge under diabetic glucose conditions, and several clocks also distinguished senescence from proliferative stages of the cellular lifespan. In gene-based analyses, both DNAm topology across genes and age-related changes were highly conserved with the *in vivo* age-related DNAm patterns across several genes, including *ELOVL2*. Finally, this study provides proof-of-concept of the experimental utility of high temporal density of DNAm measures, revealing a broad range of age-associated DNAm profiles at the single CpG level. These kinetic profiles include linear, quadratic and cubic trajectories whose biological basis and functional significance remain to be established. The conserved and accelerated nature of epigenetic aging in this cultured fibroblast system opens the door for future studies to examine how age-related DNAm trajectories are influenced by genomic and environmental factors, and whether they can predict health outcomes.

Previous *in vivo* human and mouse studies have reported an overall global DNA hypomethylation with age. More recent and sensitive studies using next-generation sequencing have provided a more nuanced picture (Unnikrishnan et al., 2018), yet the global methylation of specific cell types, including senescent cells are still thought to decrease with age. Similarly we found that hypomethylation was both overrepresented and occurred at a greater rate during cellular aging in cultured human fibroblasts. This effect was observed in CpGs globally (−2% changes in mean methylation across all probes), and through an overrepresentation of CpGs undergoing hypomethylation defined either by a lenient cutoff based on rate of methylation (77%, >5% per month), and amongst CpGs with a stringent cutoff based upon significance (79%, 1,000 lowest p values, p < 5.98e-45). These data align with those of Cruickshanks et al. (Cruickshanks et al., 2013), who found by whole-genome single-nucleotide bisulfite sequencing a 7% decrease in global methylation in senescence human lung fibroblasts (IMR90) compared to younger cells. Furthermore, regional analyses revealed that hypomethylated sites tend to map to unclassified intergenic regions and open seas, and are less commonly found in CpG islands and gene promoters. These findings suggest that rather than regulating transcriptional activity, age-related hypomethylation may influence other aspects of the genome, such as genomic stability (Benayoun, Pollina, & Brunet, 2015; Hu, Li, & Duan, 2018).

All four DNAmAge clocks successfully predicted an increase in epigenetic age correlating to various extent with the amount of time cells were grown in culture. Clocks trained on non-fibroblast tissues showed linear increases in the early and mid-life, but not during replicative senescence, while the Skin & Blood clock linearly captured aging throughout the lifespan, including during senescence. This divergence in clock capability likely stems from the source data on which the clocks were trained. The Skin & Blood clock was trained on fibroblasts at various ages, including senescent cells, which would allow the elastic-net regression models underlying the clock’s algorithm to select CpG sites that change continually through all cellular aging phases, independently of cellular division. More importantly, because the Skin & Blood clock accurately captures DNAm aging signatures in human tissues, the successful linear prediction of age *in vitro* highlights the conservation of an epigenetic aging signature between *in vivo* and *in vitro* systems.

Moreover, because the clocks provide accurate estimates of biological aging that predict one year of DNAm age per calendar year in human tissues (r>0.95), employing this system in cultured cells allowed us to the directly calculate and compare the rate of aging *in vitro*. *In vivo*, algorithms are trained to have a slope of 1. The same clocks applied to cultured fibroblasts return an average slope of about 62 DNAmAge years per calendar year, indicating that the rate of biological aging in culture is substantially faster than the aging process in the human body. Our confidence in these estimates are reinforced by the high temporal density of measures, approximately every 11 days, taken across the cellular lifespan. Both this repeated-measures design and the application of precise DNAm clocks trained on human tissues provide the necessary combination of approaches to measure the rate of aging in cellular cultures.

The basis for the observed accelerated aging remains to be established but three lines of evidence indicate that *in vitro* age acceleration is not solely due to a higher division rate of cultured cells. First, the Skin & Blood clock linearly predict an increase in DNAmAge even during replicative senescence when cells are undergoing minimal divisions. This finding aligns with recent work by Kabacik et al. (2018) showing that DNAmAge continues to increase indefinitely if cultured cells are immortalized by overexpressing the telomerase reverse transcriptase hTERT. Second, it is possible to reset epigenetic age to zero in induced pluripotent stem cells, even after extensive cell divisions (Horvath, 2013; Olova, Simpson, Marioni, & Chandra, 2019), thus uncoupling association between division and DNAmAge. Third, in human tissues epigenetic clocks accurately predict increasing DNAm age with good precision in a range of cell types that vary in their division rates by several orders of magnitudes - such as fast dividing white blood cells, and non-dividing brain neurons (Horvath, 2013). Combined, these studies suggest that the accelerated aging process captured by DNAmAge in culture can not be explained solely by a faster rate of cellular division *in vitro*.

Moreover, because we directly monitored the number of cell divisions in cultured cells and validated the use of the clocks in this system, it is possible to extrapolate the number of cell divisions per DNAmAge. This yields an estimated rate of cell division of ∼3.0 cell divisions/year. Quantifying the rate of cell division or the lifespan of different cells in the human body is challenging (Spalding, Bhardwaj, Buchholz, Druid, & Frisén, 2005). Although our approach represents the closest estimation of fibroblast division rate yet reported, this estimate is derived exclusively from the proliferative portion of the cellular lifespan where cell division rate is stable, it reflects cell populations rather than single cells, and several other factors could influence this *in vitro*-to-*in vivo* conversion, suggesting caution in the interpretation of this metric.

There is a need to be able to examine biological functions across human lifespan for two main reasons. First, aging-modifying interventions must be tested across the lifespan. But this is challenging in long-lived humans, which age at a rate too slow enable life-long longitudinal studies of individuals yet. Second, there is the need to test the influence of disease-causing and lifespan-shortening exposures – such as environmental toxicants, chronic psychosocial stress, chronic hyperglycemia. The general scientific approach has mostly relied on model organisms with shorter lifespan, such as rodents, fish, flies, or worms (Mitchell, Scheibye-Knudsen, Longo, & de Cabo, 2015). But there are limitations associated with the study of non-human organisms, which exhibit several genetic and biochemical features not shared with humans and may yield results that ultimately do not translate to human populations (de Magalhães, 2014; Hunter, 2008). The current study presents a potential solution to overcome these limitations. Primary fibroblasts are grown under controlled laboratory conditions where exposures can be precisely controlled both in time and magnitude, and exhibit DNAmAge signatures reflecting decades of biological aging within just a few months. These conditions could allow for rapid testing of age-modifying interventions with frequent longitudinal measurements in a human genetic system. For example, metabolic stress, which results in obesity and metabolic syndrome with shortened lifespan, has been associated with accelerated epigenetic aging in the liver, with an average 3.3 years of increased epigenetic age for every 10 body mass index (BMI) units (Horvath et al., 2014). Here, chronic diabetic glucose levels caused a stepwise increase in DNAmAge of ∼2.9 years in fibroblasts, without changing the rate of aging. This form of age acceleration previously referred to as epigenetic age acceleration (EAA) (Chen et al., 2016) reflects an acute shifts in age position (i.e., intercept of linear function) and should be distinguished from changes in the rate of aging (i.e., the slope) affecting lifelong trajectories. In future studies, this experimental system could be used to predict the effect of metabolic or chemical stressors, or the effectiveness of senolytic and therapeutic interventions on human aging trajectories over a shortened time scale. It also remains to be determined how much the rate of aging varies between individuals, and whether it is predictive of future health outcomes.

Three independent findings indicated that cellular aging *in vitro* and *in vivo* are conserved biological process: (i) DNAmAge clocks trained *in vivo* track aging *in vitro*; (ii) both systems show global hypomethylation across lifespan, (iii) cell replication-based methylation errors (MiAge) are conserved. We then further examined the likelihood that *in vitro* cellular aging follows the same patterns as tissue-based human aging signatures through a candidate gene approach focusing on *ELOVL2* and its associated promoter CpG cg16867657, previously validated across multiple cohort and longitudinal studies from human blood (Wang et al., 2018). Under most circumstances, DNAm of promoter-associated CpG islands generally decreases the expression of its downstream gene through recruitment of repressor elements (Schübeler, 2015). In relation to *ELOVL2*, we found that the topology of DNAm across the entire gene was highly conserved, and the age-related hypermethylation of cg16867657. The linear hypermethylation of cg16867657 was associated with robust downregulation of ELOVL2 transcript levels, consistent with the functional significance of this age-related DNAm change. Specifically, a 25% increase in methylation levels in cg16867657 was associated with an 83% decreased expression over a period corresponding to ∼10 years of human life. In contrast to previous studies in human tissues, this level DNAm-to-gene expression correlation is remarkably strong, possibly owing to the influence on environmental factors on *ELOVL2* expression in humans, compared to the absence of these factors *in vitro*, which may reduce noise and facilitate the detection of meaningful genomic-epigenomic interactions.

In relation to lifespan kinetics at single CpG resolution, a notable finding is the non-linear behavior of hypomethylated and hypermethylated CpGs. In longitudinal human aging studies, researchers often rely on linear models to identify age-related associations, and have access to a limited number of timepoints stemming from the considerable challenge that following participants over decades represents (Alisch et al., 2012; N. D. Johnson et al., 2017). Here using high-temporal resolution data, we highlight the variety of aging kinetics in the methylome across the cellular lifespan. Whereas the vast majority of CpGs did not change with age, our analysis indicates that of the sites that do undergo hypo- or hypermethylation, most (>90%) behave non-linearly, with their age-related patterns being best described with 2 to 4 degrees of freedom. As expected, we also found that most CpG methylation trajectories are monotone functions of time (increasing or decreasing), although some CpGs may exhibit non-monotone trajectories (e.g., increase during early life, then decrease during mid-life and senescence). With twelve timepoints across age we could detect up to three directional inflection points over time, and more frequent measurements could allow to discover with still greater confidence more complex age-related kinetics. The multi-directional changes in DNA methylation in some sites is consistent with previously described short-lived oscillations in DNAm (circadian and seasonal) (Lim et al., 2017, 2014; Oh et al., 2018, 2019). These data highlight the value model systems that enable high-temporal resolution sampling and statistical models that accommodate non-linear DNAm dynamics.

Overall, this proof-of-concept work establishes the conserved yet accelerated DNAm aging of cultured fibroblasts at the global, gene, and single CpG levels. Using this system and its high-temporal resolution, we propose that it is possible to leverage both integrative clocks that capture human aging signatures, as well as non-linear modeling methods to capture the full spectrum of age-related DNAm changes. Although our work suggests that aging signatures are largely conserved between whole blood and cultured fibroblasts, future work is needed to fully define the epigenomic overlap among various human tissues and cell types (Slieker, Relton, Gaunt, Slagboom, & Heijmans, 2018) and across individuals. Longitudinally quantifying the *rate* of aging in primary human cells may also enable to rapidly assess the effectiveness of age-modifying interventions across the lifespan.

## Experimental Procedures

The detailed version of the Experimental Procedures is available in *Appendix 1*.

### Cell culture

Primary human dermal fibroblasts were obtained with informed consent from a healthy 29-year-old male donor (IRB #AAAB0483). Cells were maintained in standard 5% CO_2_ and atmospheric O_2_ at 37°C in T75 flasks. Cells were passaged approximately every five days, which corresponded to ∼90% confluency, with 500,000 cells replated each passage. Brightfield microscopy images (20x magnification) were taken using inverted phase-contrast microscope. The starting passage numbers was three, and cells were terminated after exhibiting less than one population doubling over a 30-day period.

### DNA methylation sample preparation

Twelve normal glucose (5.5mM) timepoints and fourteen high glucose (25mM) timepoints were collected approximately every 11 days were selected for DNA Methylation measurements. DNA was extracted using a DNeasy kit (Qiagen cat#69506) according to the manufacturer’s protocol. At least 150 ng of DNA was submitted in 50 µl of ddH2O to the New York Genome Center for bisulfite conversion and hybridization using the Infinium Methylation EPIC BeadChip kit. Samples were randomly distributed across two plates. DNA Methylation levels were measured for 866,836 CpG Sites.

## Data preprocessing and quality control

All DNA methylation data was processed in R (Version 3.5.0). Quality control preprocessing was applied by checking for correct sex prediction, probe quality, sample intensities, and excluding SNPs and non-CpG probes. All samples passed our quality control and no samples were excluded. Data was then normalized using Functional Normalization. Using the R package SVA, both RCP and ComBat adjustments were applied for probe-type and plate bias, respectively. These adjustments excluded 68 out of the 866,836 CpG probes for a final of 866,768 probes for further analysis. On average, probes had a coefficient of variation of 10.08% with a bias in technical error towards demethylated sites (Figure S1B).

### PCA analysis

Principal Component Analysis was performed using ‘prcomp’ with zero centering and unit scaling. PCA was applied on normal glucose samples, high glucose samples, and then on all samples together. All three analyses showed similar components and age-related effects regardless of glucose treatment or number of samples included.

### Rate of methylation change

For global rates of methylation change over time, we applied a linear regression model to each CpG site across the lifespan and then transformed the slope to percent change per month. Sites were designated as hyper- or hypo- methylated with age based of the sign of the regression slope, as plotted in Figure 1.

### DNAmAge clocks and quantification of the rate of DNAm aging

The four DNAmAge clocks predicted epigenetic age was predicted using linear coefficients detailed in the respective clock’s source paper (Appendix S1 Table 1).

For the Hannum clock, six of 71 clock sites are not present in the EPIC array used in our study. These missing sites explain why the reported DNAmAge is not in recognizable units of time. The rate of aging is determined by the first derivative (i.e., slope) of the linear regression between chronological time and DNAmAge. All algorithms are trained to accurately predict age and thus have a slope of 1 in human tissues. To assess age-acceleration, we compared the measured slope of DNAmAge across lifespan and compared it to the theoretical value of 1, or to other slopes (normal vs high glucose). Stepwise age-increase is defined as increase in the y intercept of DNAmAge across the lifespan without a change in slope.

### Mitotic Age calculator

MiAge estimations were calculated using the previously described calculator (Youn & Wang, 2018). Briefly, using the stochastic replication errors accumulated in the epigenetic inheritance process during cell divisions, the MiAge calculator was trained on 4,020 tumor and adjacent normal tissue samples from eight TCGA cancer types, which consists of a panel of 268 CpGs together with their estimated site-specific parameters.

### GAM model and single site analysis

To assess the range of kinetic complexity across the cellular lifespan, we fitted each of the 866,736 CpGs with generalized additive models (GAM) using the MGCV package. We then used each GAM model to estimate the change in methylation (beta values) and the rate of change across the cellular lifespan. The top 1,000 most significant CpGs were selected and classified according to their degree of freedom (DoF). The maximum DoF determined to be biologically meaningful was 4, as increasing DoF up to 10 did not identify additional significant CpG hypo- or hyper-methylation trajectories. We then assessed the prevalence of CpGs at different genomic loci grouped based on their estimated DoF. CpGs were annotated with gene location and related regulatory features using the manufacturer’s (Illumina EPIC array) annotation.

### In vivo whole blood data

The longitudinal *in vivo* DNA methylation data were from Swedish Adoption/Twin Study of Aging (SATSA), part of the Swedish Twin Registry, which is a national register of twins born between 1886-2000, as previously described in details in Wang et al. (2018). This *in vivo* dataset contains DNA methylation levels measured using the 450k array on whole blood DNA from 385 participants (including 85 monozygotic and 116 dizygotic twin pairs). The longitudinal component included up to 5 timepoints approximately every 3 years, for a total of 1,011 samples after quality control. Data was processed as described previously (Wang et al. 2018) and both gene DNAm topology and single-CpG trajectories were compared with data from human fibroblasts.

### In vivo fibroblast data

Age-related CpGs of *In Vivo* fibroblasts were obtained from Horvath et al. 2018. Briefly, fibroblasts were collected from 147 donors ranging from 0 to 100 years old and DNAm measurements were obtained using Illumina 450k array. 28,744 age-related CpGs were identified using Standard screening for numeric traits with a biweight midcorrelation using the R package WGCNA v1.66 and corrected for multiple comparisons using bonferroni correction.

### Gene-based DNA methylation topology

Age-related CpGs were selected from the top 29 genes identified by Wang et al. 2018 using mixed-effects models across *in vivo* lifespan. The overlap of *in vitro* and *in vivo* methylation for individual genes was assessed using the EPIC Illumina array annotation. After selecting CpG sites of a given gene, sites were ordered by gene position and then annotated with related related-gene regions and CpG islands. The positions of CpGs were visualized using the UCSC genome browser (https://genome.ucsc.edu/).

### RNAseq

Total genomic RNA was isolated at three timepoints across the cellular lifespan. At each timepoint, ∼2 million cells were stored in 1ml trysol (Invitrogen cat#15596026), RNA was extracted on-column DNAse treated according to the manufacturer’s instructions, and quantified using the QUBIT high sensitivity kit (Thermo Fisher Scientific cat#Q32852). RNA was submitted to the Columbia Genome Center and sequenced (Illumina NovaSeq 6000), yielding approximately 30 million 100 bp single-end reads. Transcript levels are shown as fragments per kilobase of transcript per million mapped reads (FPKM) expressed relative to the youngest timepoint.

## Acknowledgements

Work of the authors is supported by NIH grants GM119793, MH113011, MH119336 (M.P.); OD021412 (M.A.); AG060908 (S.Horvath); HD32062, NS078059; LM013061 (S.W.); and CA226672 (M.H.). The authors are also supported by the Wharton Fund and the Columbia Aging Center (M.P.), the John Harvard Distinguished Science Fellow Program within the FAS Division of Science of Harvard University (M.A.); and the J. Willard and Alice S. Marriott Foundation, Muscular Dystrophy Association, Arturo Estopinan TK2 Research Fund, Nicholas Nunno Foundation, JDF Fund for Mitochondrial Research, and Shuman Mitochondrial Disease Fund (M.H.). The SATSA study is supported by NIH grants AG04563; AG10175; AG028555, the MacArthur Foundation Research Network on Successful Aging, the European Union’s Horizon 2020 research and innovation programme No. 634821, the Swedish Council for Working Life and Social Research (FAS/FORTE) 97:0147:1B, 2009-0795, 2013-2292, the Swedish Research Council 825-2007-7460, 825-2009-6141, 521-2013-8689, 2015-03255, 2015-06796, the Karolinska Institutet delfinansiering (KID) grant for doctoral students (Y.W.), the KI Foundation, the Strategic Research Area in Epidemiology at Karolinska Institutet and by Erik Rönnbergs donation for scientific studies in aging and age-related diseases. The content is solely the responsibility of the authors and does not necessarily represent the official views of the National Institutes of Health.

## Author Contributions

G.S. and M.P. designed experiments. G.S. performed tissue culture experiments and collected DNA methylation data. G.S., A.C., and M.A.B. processed and analyzed *in vitro* DNA methylation data. S.W., S.Horvath, S.Hägg, and Y.W. provided *in vivo* methylation data and clock algorithms. M.H. provided cells. G.S. and M.P. drafted manuscript and prepared figures. All authors edited and approved the final version of the manuscript.

## Supporting Information Listing

**Appendix S1.** Full length version of Experimental Procedures.

**Table S1.** Overlapping CpGs across in vivo in vitro whole blood and fibroblast datasets

**Figure S1**. Comparison of EPIC array normalization methods and quantification of technical variability.

**Figure S2**. The PhenoAge clock, Hannum clock, and MiAge calculator track cellular aging in early- and mid-life.

**Figure S3**. DNAm gene topology and age-related CpG differences are generally conserved between human blood and cultured fibroblasts.

**Figure S4**. Overlap of age-related DNA methylation changes between in vitro and in vivo fibroblasts and whole blood.

**Figure S5**. Exploratory analysis of non-monotone CpG trajectories.

**Figure S6**. Age-related DNAm gene topology graphs illustrate the degree of conservation between human blood and cultured fibroblasts.

