## Supplemental_Methods for "Human Aging DNA Methylation Signatures are Conserved but Accelerated in Cultured Fibroblasts"

**Appendix S1. Full length version of Experimental Procedures.**

***Cell culture***

Primary human dermal fibroblasts were obtained with informed consent from a healthy 29-year-old male donor (IRB #AAAB0483). Cryopreserved cells were seeded cultured in Dulbecco's modified Eagle's medium (DMEM, Thermo Fisher #10569044) containing, 1 mM pyruvate, 4 mM GlutaMAX$^{TM}$, supplemented with 10% fetal bovine serum (FBS; not heat inactivated), 1% non-essential amino acids (NEAA), and either normal (5.5mM) or high (25mM) glucose. After initial plating cells were frozen using cryo-media (10% DMSO in DMEM) in liquid nitrogen and later thawed for experimentation. Cells were maintained in standard 5% $CO_2$ and atmospheric $O_2$ at 37°C in T75 flasks. Cells were passaged approximately every five days, which corresponded to ~90% confluency, with 500,000 cells replated each passage. As cell growth slowed the number of days between passages was also continually extended to maintain 90% confluency at the point passage. At each passage, cells were counted using the Countess II Automated Cell Counter (Thermo Fisher Scientific cat#AMQAX1000), and the counts used to compute population doublings. Brightfield microscopy images (20x magnification) were taken using inverted phase-contrast microscope. The starting passage numbers was three, and cells were terminated after exhibiting less than one population doubling over a 30-day period.

***DNA methylation sample preparation***

Twelve normal glucose (5.5mM) time points and fourteen high glucose (25mM) time points were collected approximately every 11 days were selected for DNA Methylation measurements. DNA was extracted using a DNeasy kit (Qiagen cat#69506) according to the manufacturer's protocol. At least 150 ng of DNA was submitted in 50 µl of ddH2O to the New York Genome Center for bisulfite conversion and hybridization using the Infinium Methylation EPIC BeadChip kit. Samples were randomly distributed across two plates. DNA Methylation levels were measured for 866,836 CpG Sites.

### Data preprocessing and quality control

All DNA methylation data was processed in R (Version 3.5.0). Quality control preprocessing was applied by checking for correct sex prediction, probe quality, sample intensities, and excluding SNPs and non-CpG probes. All samples passed our quality control and no samples were excluded. Data was then normalized using Functional Normalization. Several other normalization methods were tested on the EPIC chip including Noob, SWAN, Illumina, and Quantile to evaluate its impact on DNAmAge calculations (Figure S1A). Regardless of normalization method used, the relationship of the Pan-Tissue DNAmAge with cellular age was maintained within a $r^2$ between 0.88-0.92. Using the R package SVA, both RCP and ComBat adjustments were applied for probe-type and plate bias, respectively. These adjustments excluded 68 out of the 866,836 CpG probes for a final of 866,768 probes for further analysis. On average, probes had a coefficient of variation of 10.08% with a bias in technical error towards demethylated sites (Figure S1B).

#### *PCA analysis*

Principal Component Analysis was performed using 'prcomp' with zero centering and unit scaling. PCA was applied on normal glucose samples, high glucose samples, and then on all samples together. All three analyses showed similar components and age-related effects regardless of glucose treatment or number of samples included.

#### *Rate of methylation change*

For global rates of methylation change over time, we applied a linear regression model to each CpG site across the lifespan and then transformed the slope to percent change per month. Sites were designated as hyper- or hypo- methylated with age based of the sign of the regression slope, as plotted in Figure 1.

#### *DNAmAge clocks and quantification of the rate of DNAm aging*

The four DNAmAge clocks predicted epigenetic age was predicted using linear coefficients detailed in the respective clock's source paper (Appendix S1 Table 1).

| Clock | Source Paper | Learning Algorithm | CpGs | Trained Tissue Type |
|---|---|---|---|---|
| Pan-Tissue | Horvath 2013, Genome Biology PMID: 24138928 | Elastic-Net Regression | 353 | 51 Tissue Types including several tumor types |
| Skin & Blood | Horvath 2018, Aging PMID:30048243 | Elastic-Net Regression | 391 | Fibroblasts, keratinocytes, buccal cells, endothelial cells, lymphoblastoid cells, skin, blood, and saliva |
| PhenoAge | Levine 2018, Aging PMID: 29676998 | Elastic-Net Regression | 513 | Whole Blood |
| Hannum | Hannum 2013, Molecular Cell PMID: 23177740 | Elastic-Net Regression | 71 | Whole Blood |

**Appendix S1 Table 1. DNAmAge Clocks Descriptions.**

Table of the Pan-Tissue, Skin & Blood, PhenoAge, and Hannum clock descriptions.

For the Hannum clock, six of 71 clock sites are not present in the EPIC array used in our study. These missing sites explain why the reported DNAmAge is not in recognizable units of time. The rate of aging is determined by the first derivative (i.e., slope) of the linear regression between chronological time and DNAmAge. All algorithms are trained to accurately predict age and thus have a slope of 1 in human tissues. To assess age-acceleration, we compared the measured slope of DNAmAge across lifespan and compared it to the theoretical value of 1, or to other slopes (normal vs high glucose). Stepwise age-increase is defined as increase in the y intercept of DNAmAge across the lifespan without a change in slope.

***Mitotic Age calculator***

MiAge estimations were calculated using the previously described calculator (Youn & Wang, 2018). Briefly, using the stochastic replication errors accumulated in the epigenetic inheritance process during cell divisions, the MiAge calculator was trained on 4,020 tumor and adjacent normal tissue samples from eight TCGA cancer types, which consists of a panel of 268 CpGs together with their estimated site-specific parameters.

#### *GAM model and single site analysis*

To assess the range of kinetic complexity across cellular lifespan, we fit 866,736 generalized additive models (GAM) using the MGCV package. The GAM model uses regularized, nonparametric functions (penalized regression splines) that allow for more fitting flexibility because the relationships between independent and dependent variable can be non-linear. Further, the GAM model outputs the contribution of each independent variable to the prediction allowing for high interpretability. This includes the estimated degrees of freedom (DoF), which corresponds to the smoothness of the fitted function. The GAM model estimates the change in methylation (beta values) across cellular lifespan. The top thousand most significant CpGs were selected and classified according to their DoF. The maximum degree of freedom determined to be biologically meaningful was 4, as increasing DoF up to 10 did not identify additional significant CpG hypo- or hyper-methylation trajectories. We then assessed the prevalence of CpGs at different genomic loci grouped based on their estimated DoF. CpGs were annotated with gene location and related regulatory features using the manufacturer's (Illumina EPIC array) annotation.

#### *In vivo whole blood data*

The longitudinal *in vivo* DNA methylation data were from Swedish Adoption/Twin Study of Aging (SATSA), part of the Swedish Twin Registry, which is a national register of twins born between 1886-2000, as previously described in details in Wang et al. (2018). This dataset can be found at the Arrayexpress database of EMBL-EBL (www.ebi.ac.uk/arrayexpress) under accession number E-MTAB-7309. This *in vivo* dataset contains DNA methylation levels measured using the 450k array on whole blood DNA from 385 participants (including 85 monozygotic and 116 dizygotic twin pairs). The longitudinal component included up to 5 time points approximately every 3 years, for a total of 1011 samples after quality control. Data was processed as described previously (Wang et al. 2018) and both gene DNAm topology and single-CpG trajectories were compared with data from human fibroblasts.

#### *In vivo fibroblast data*

Age-related CpGs of *In Vivo* fibroblasts were obtained from Horvath et al. 2018. Briefly, fibroblasts were collected from 147 donors ranging from 0 to 100 years old and DNAm measurements were obtained using Illumina 450k array. 28,744 age-related CpGs were identified using Standard screening for numeric traits with a biweight midcorrelation using the R package WGCNA v1.66 and corrected for multiple comparisons using bonferroni correction.

#### *Gene-based DNA methylation topology*

Age-related CpGs were selected from the top 29 genes identified by Wang et al. 2018 using mixed-effects models across *in vivo* lifespan. The overlap of *in vitro* and *in vivo* methylation for individual genes was assessed using the EPIC Illumina array annotation. After selecting CpG sites of a given gene, sites were ordered by gene position and then annotated with related related-gene regions and CpG islands. Only sites appearing on both 450k and EPIC chips were included in topology graphs. Additionally, if genes had more than 30 CpG sites, topology graphs were trimmed to 15 sites in both directions around the gene's most age-related CpG. Correlational matrices were composed of all CpG sites regardless of the number of CpG sites on the gene. The positions of CpGs were visualized using the UCSC genome browser (https://genome.ucsc.edu/).

#### *RNAseq*

Total genomic RNA was isolated at three timepoints across cellular lifespan. At each timepoint, ~2 million cells were stored in 1ml trysol (Invitrogen cat#15596026), RNA was extracted on-column DNAse treated according to the manufacturer's instructions, and quantified using the QUBIT high sensitivity kit (Thermo Fisher Scientific cat#Q32852). RNA integrity and concentration was also assessed by bioanalyzer 2100 (Agilent); all samples had RNA integrity numbers (RIN) > 9. A total of 500 ng of RNA was submitted in 50 μl of RNAase free water to the Columbia Genome Center. Ribosomal RNA was removed using ribo-zero gold (Illumina cat#MRZE706), cDNA libraries were prepared using the TrueSeq Stranded Total RNA Library Prep Kit (Illumina cat#20020596), and sequenced (Illumina NovaSeq 6000), yielding approximately 30 million 100 bp single-end reads. Base calls were performed using RTA (Illumina) and then converted to fastq format using bclfastq2 (version 2.20). Reads were then mapped to a reference genome (Human: NCBI/build37.2) using STAR (2.5.2b) and featureCounts (v1.5.0-p3). RNA levels were quantified as fragments per kilobase of transcript per million mapped reads (FPKM). Transcript levels are shown as FPKM values expressed relative to the youngest timepoint.
