## Supplemental_Figures for "Human Aging DNA Methylation Signatures are Conserved but Accelerated in Cultured Fibroblasts"

| Count | CpG Site | UCSC Gene | UCSC Region | Relation to Island | Methylation (Cultured Fibroblasts) | Gene Expression (Culture Fibroblasts) |
|---|---|---|---|---|---|---|
| 1 | cg12432010 | IQSEC3 | 1stExon | Island | Hyper (10%) | Undetected |
| **2** | **cg25391162** | **FOXD4** | **1stExon** | **Island** | **Hyper (15%)** | **Upregulated (333%)** |
| 3 | cg01503065 | DCHS2 | 1stExon | Island | Hyper (9%) | Not Significant |
| 4 | cg00059225 | GLRA1 | 1stExon;5'UTR | Island | Hyper (14%) | Undetected |
| 5 | cg06665453 | TSEN54;LLGL2 | 3'UTR;TSS1500 | N_Shore | Hyper (18%) | Not Signifcant;Undetected |
| 6 | cg00741624 | KIAA1409 | 5'UTR | Island | Hyper (8%) | Undetected |
| 7 | cg25032595 | CLDN10 | 5'UTR;1stExon;Body | Island | Hyper (27%) | Not Significant |
| **8** | **cg17593472** | **FGF1** | **5'UTR;1stExon;Body;TSS200** | **OpenSea** | **Hyper (21%)** | **Upregulated (361%)** |
| 9 | cg15341124 | DIO3;MIR1247 | 5'UTR;1stExon;TSS1500 | Island | Hyper (7%) | Undetected;Undetected |
| 10 | cg00003345 | CASZ1 | 5'UTR | OpenSea | Hyper (21%) | Undetected |
| 11 | cg03615565 | FAM65A | Body | Island | Hyper (34%) | Not Significant |
| 12 | cg07544187 | CILP2 | Body | Island | Hyper (20%) | Not Significant |
| 13 | cg13654588 | PRLHR | Body | Island | Hyper (16%) | Undetected |
| 14 | cg15243034 | USP35 | Body | Island | Hyper (44%) | Not Significant |
| 15 | cg00481951 | SST | Body | N_Shore | Hyper (6%) | Undetected |
| **16** | **cg07022048** | **KRT7** | **Body** | **N_Shore** | **Hypo (-22%)** | **Upregulated (250%)** |
| 17 | cg04999352 | RARRES3 | Body | OpenSea | Hyper (8%) | Not Significant |
| 18 | cg22108374 | CCDC33 | Body | OpenSea | Hyper (46%) | Undetected |
| 19 | cg27401724 | ACBD4 | Body | S_Shelf | Hypo (-19%) | Not Significant |
| 20 | cg26470501 | BCL3 | Body | S_Shore | Hyper (21%) | Not Significant |
| 21 | cg23527621 | ECE2;CAMK2N2 | Body;3'UTR | Island | Hypo (-15%) | Not Significant;Not Significant |
| 22 | cg11071401 | CACNA1G | TSS1500 | Island | Hyper (37%) | Undetected |
| **23** | **cg16867657** | **ELOVL2** | **TSS1500** | **Island** | Hyper (22%) | **Downregulated (-83%)** |
| **24** | **cg22809047** | **RPL31** | **TSS1500** | **Island** | Hypo (-31%) | **Downregulated (-22%)** |
| 25 | cg27320127 | KCNK12 | TSS1500 | Island | Hyper (5%) | Undetected |
| 26 | cg24168221 | CST9 | TSS1500 | OpenSea | Hyper (38%) | Undetected |
| **27** | **cg25410668** | **RPA2** | **TSS1500** | **S_Shore** | Hyper (26%) | **Upregulated (35%)** |
| **28** | **cg22454769** | **FHL2** | **TSS200;5'UTR** | **Island** | Hyper (29%) | **Upregulated (155%)** |
| 29 | cg01490733 | Unclassified | Unclassified | Island | Hyper (19%) | Unclassified |
| 30 | cg15168727 | Unclassified | Unclassified | Island | Hyper (17%) | Unclassified |
| 31 | cg19821713 | Unclassified | Unclassified | Island | Hyper (23%) | Unclassified |
| 32 | cg16008966 | Unclassified | Unclassified | OpenSea | Hypo (-22%) | Unclassified |
| 33 | cg11793449 | Unclassified | Unclassified | S_Shelf | Hyper (11%) | Unclassified |
| 34 | cg17110586 | Unclassified | Unclassified | S_Shelf | Hyper (6%) | Unclassified |
| 35 | cg06931612 | Unclassified | Unclassified | S_Shore | Hyper (14%) | Unclassified |

**Supplemental Table S1. Overlapping CpGs across in vivo in vitro whole blood and fibroblast datasets**

Table of descriptive information of 35 overlapping age-associated sites between fibroblast and whole blood In Vivo and In Vitro datasets. Bolded CpG Sites indicate CpG sites on genes that are differentially expressed by >0.2 fold with age.

### Supplemental Figure S1

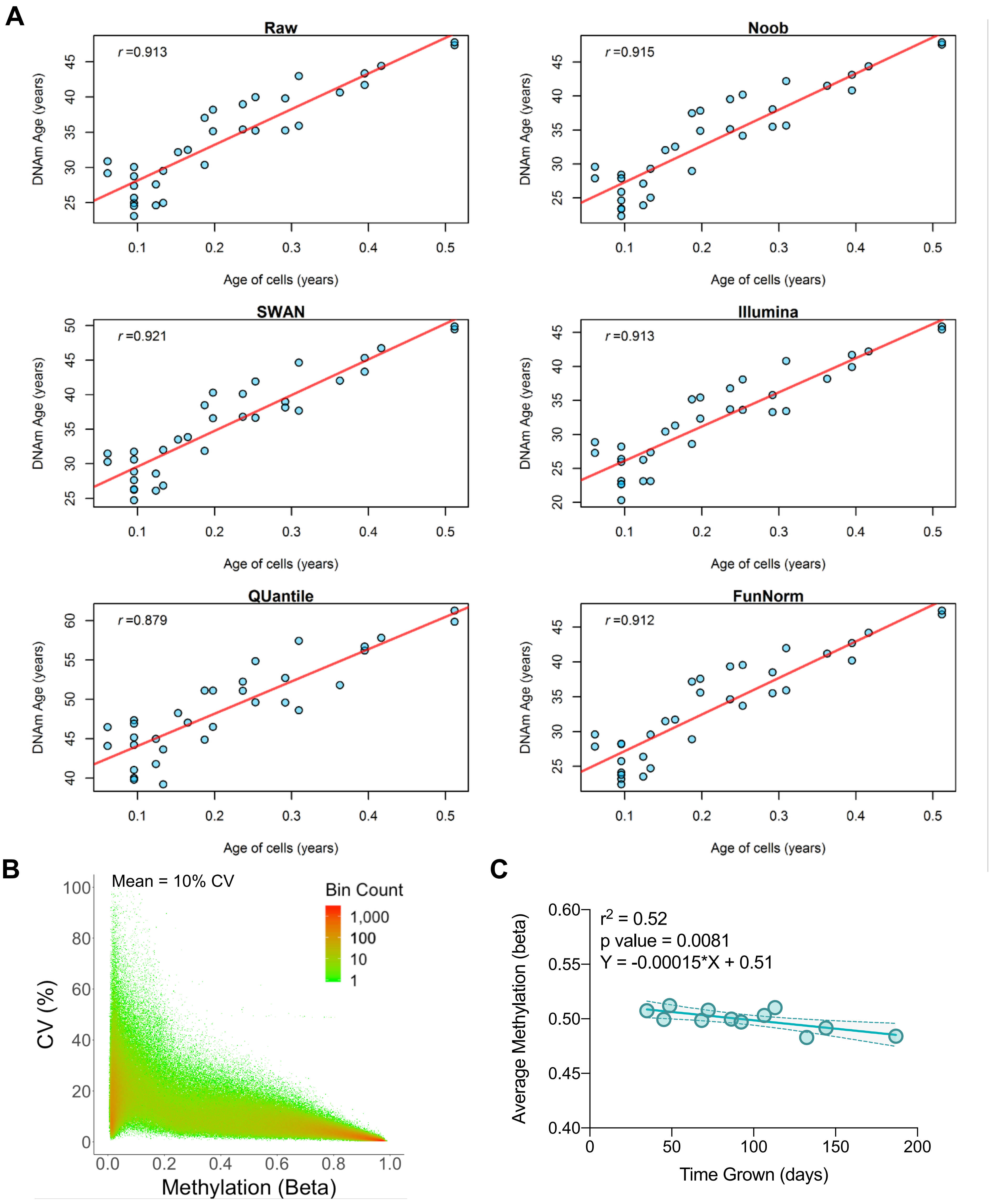


**S1. Comparison of EPIC array normalization methods and quantification of technical variability.**

(**A**) Comparisons of six normalization methods and their effect on predicted DNAmAge (PanTissue clock) regressed on chronological age. The red line indicates the linear fit line. This data demonstrates minimal effect of normalization methods on predicted DNAm age. (**B**) The calculated coefficient of variation (C.V.) measured from six technical replicates (45-day timepoint, normal glucose) distributed across different EPIC arrays. Measurement error is higher for sites with low methylation levels (beta values). Each datapoint is the C.V. for a single CpG. Color scheme indicates density of overlapping CpGs (log scale) on the scatterplot. (**C**) Linear regression of the average DNAm levels across all CpGs sites for each timepoint studied across the cellular lifespan. The global slope reflects a 5.5% decrease in global DNAm per calendar year.

### Supplemental Figure S2

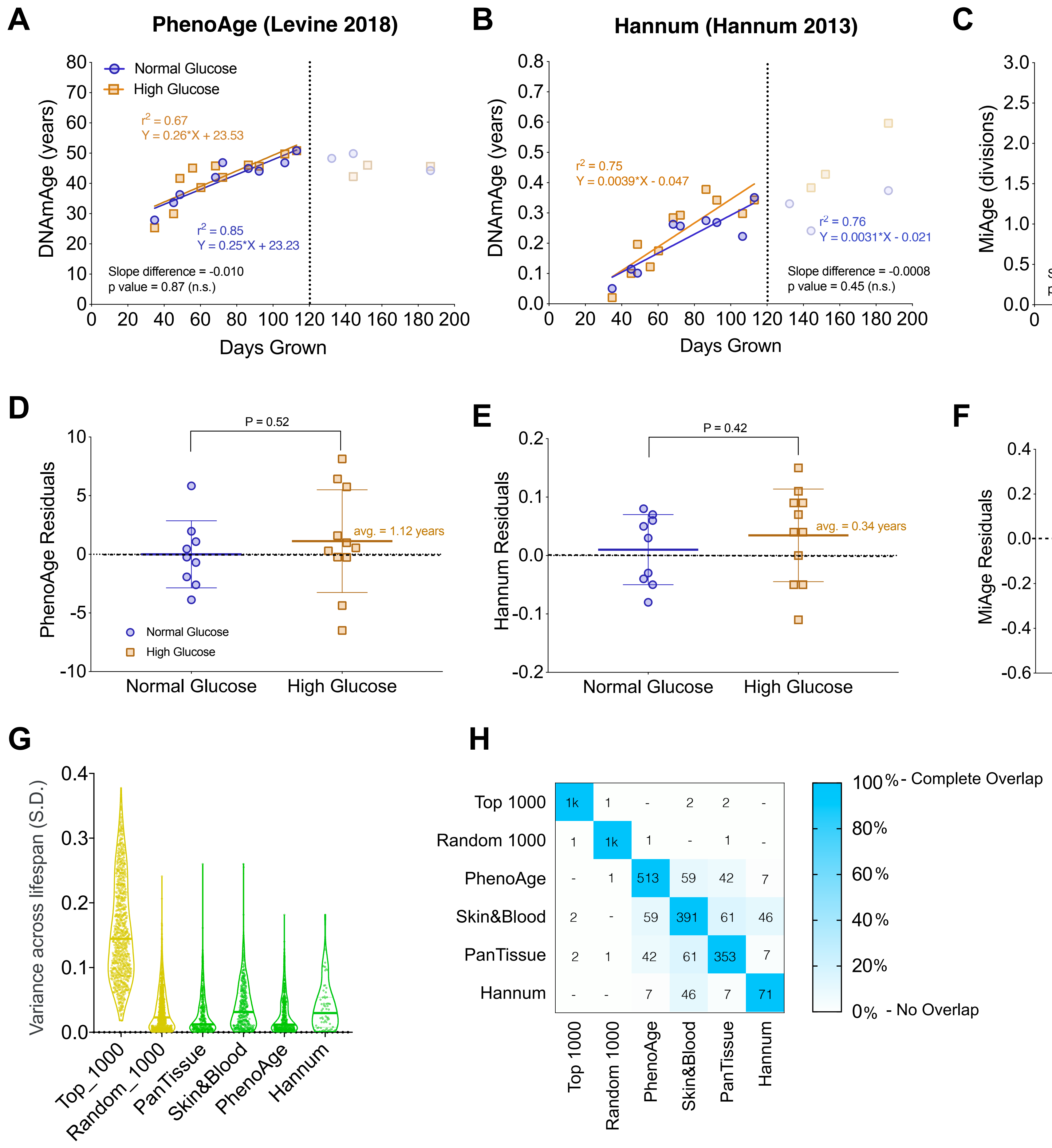


**S2. The PhenoAge clock, Hannum clock, and MiAge calculator track cellular aging in early- and mid-life.**

(**A**) The PhenoAge Clock (Levine et al., 2018) trained on blood to predict clinical age related phenotypic characteristics; (**B**) The Hannum Clock trained on blood to predict chronological age; (**C**) DNA methylation-based estimation of mitotic age trained on human tumors and blood to predict the number of cell divisions or population doublings. Shown are correlations of chronological age (days in culture) and predicted DNAmAge with each clock, indicating linear increase in DNAmAge for the early- and mid-life periods, but not beyond the point of transition towards replicative senescence (dotted line, 120 days). Results for both normal (5.5 mM) and high (25 mM) glucose are shown and do not differ in their rate of aging. (**D-F**) Box plots of residuals from the regression of actual and predicted age for normal and high glucose cells across entire lifespan. Non-parametric unpaired Mann-Whitney test. (**G**) Average age-related variation among the CpGs that compose the various DNAm clocks and the top 1,000 most significant sites in fibroblast experiments. This reveals that clock CpGs exhibit small to moderate effect sizes across the lifespan compared to the top 1,000 most significant sites in fibroblasts. All clocks show strong predictive power across the cellular lifespan despite the small effect sizes of each CpG, highlighting the added value of computationally integrating multiple CpG sites in DNAm clocks. (**H**) Proportion of overlap between different DNAmAge clocks, the top 1,000 most significant CpGs in fibroblasts, and 1,000 CpGs randomly selected from the EPIC array. Shown in each box is the number of overlapping CpGs between each group, dash indicates 0 overlapping sites.

### Supplemental Figure S3

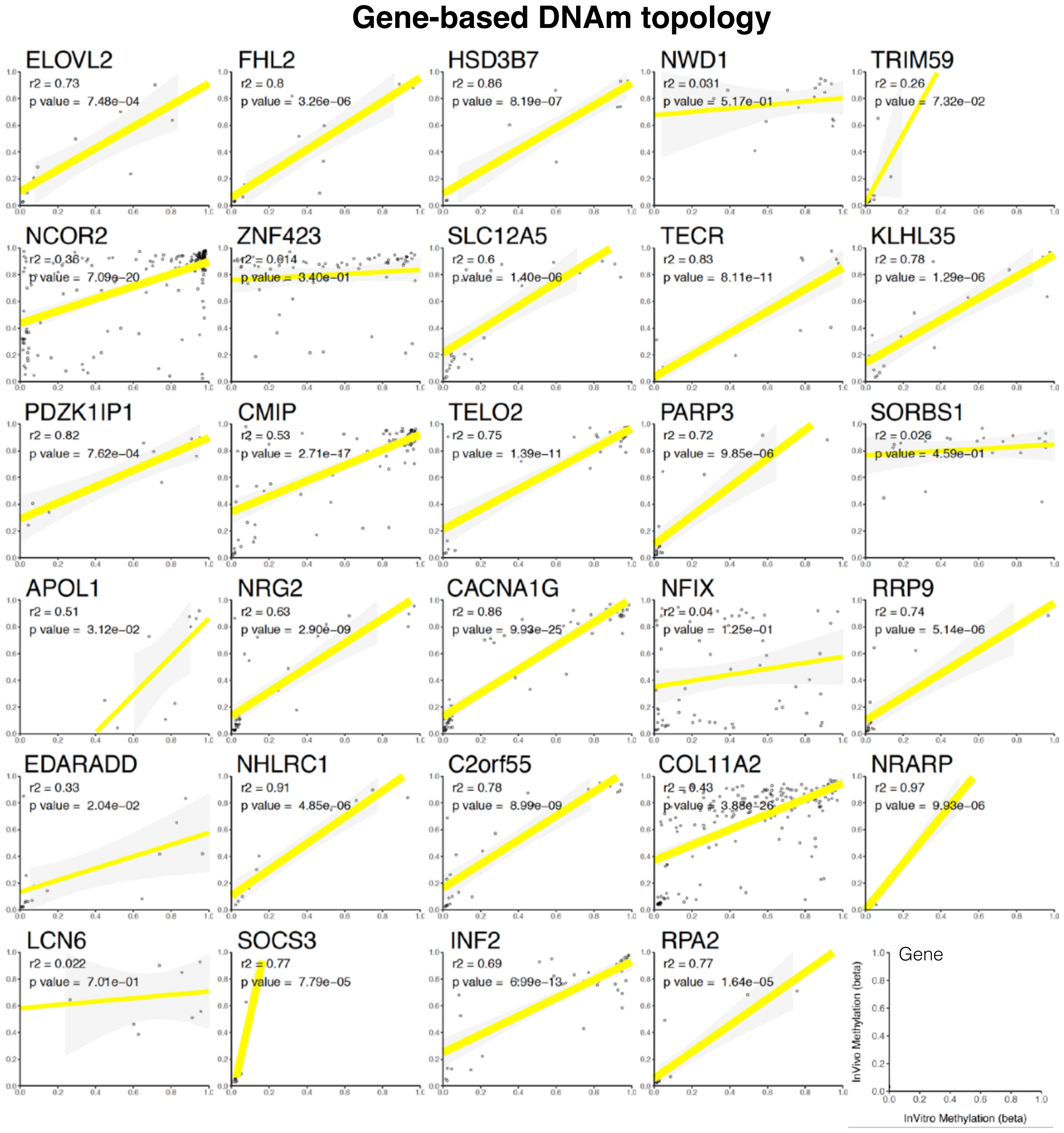


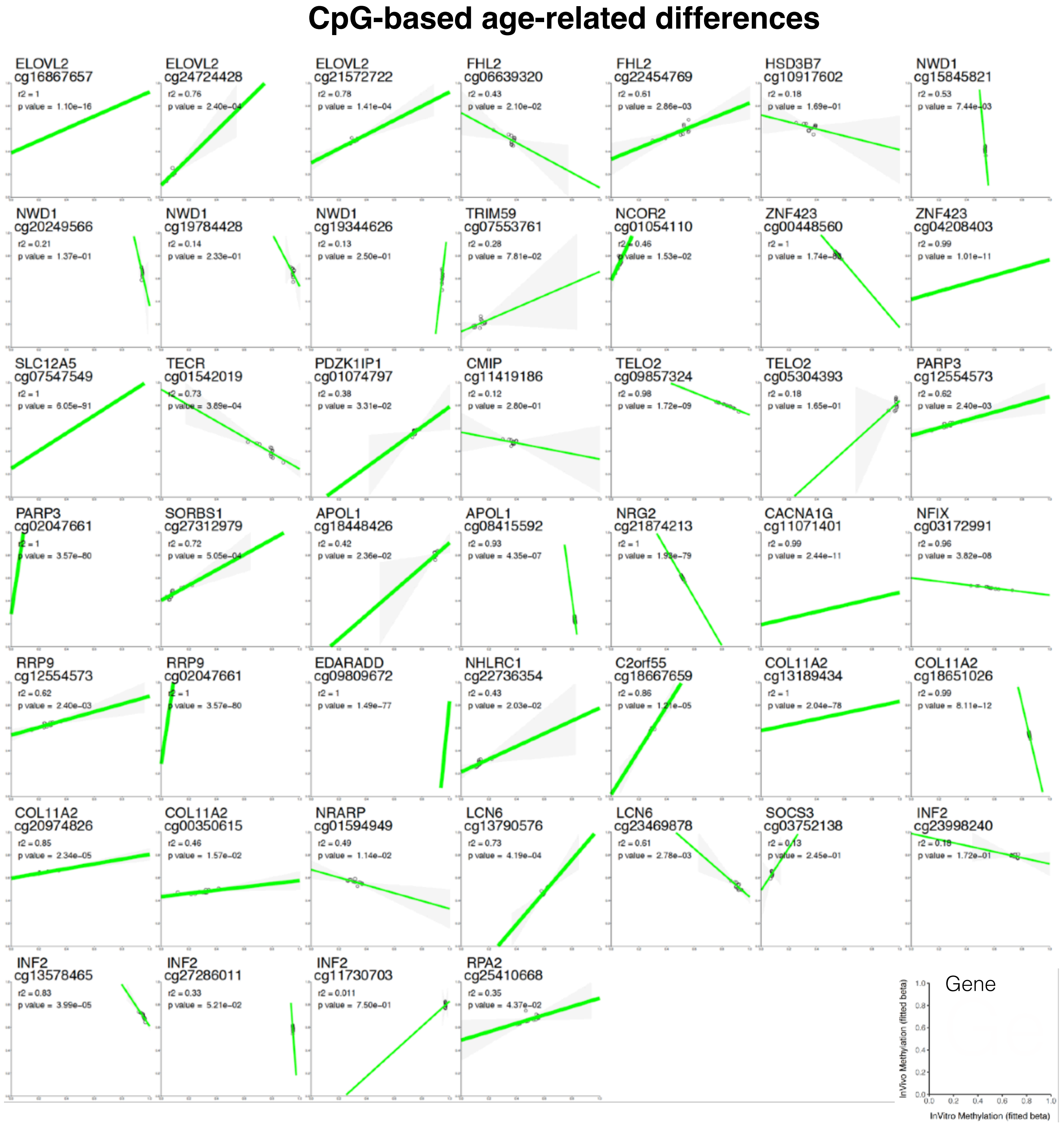


**Figure S3.** ***DNAm gene topology and age-related CpG differences are generally conserved between human blood and cultured fibroblasts.***

Higher magnification version of the Figure 3E-F, including correlation coefficients and p values. Only regressions with positive correlations are shown as significant (thick regression line).

### Supplemental Figure S4

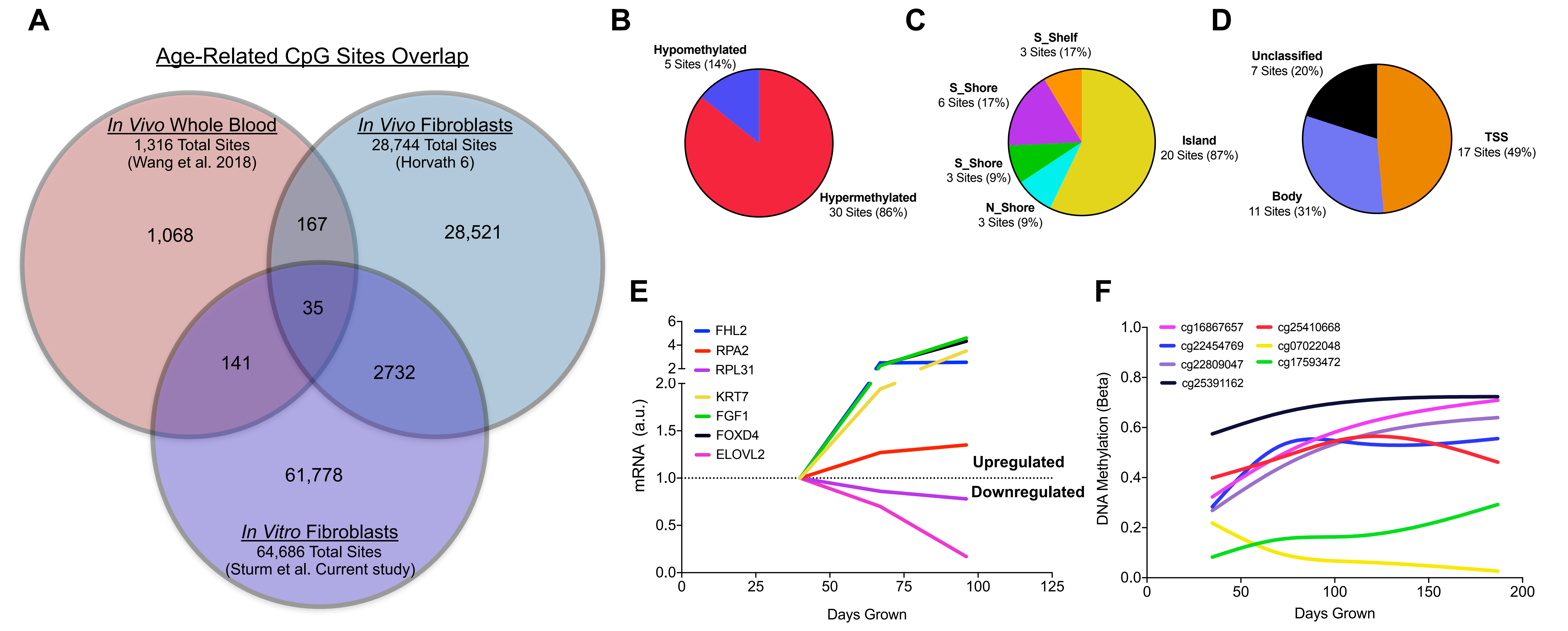


**Figure S4. Overlap of age-related DNA methylation changes between in vitro and in vivo fibroblasts and whole blood.**

(**A**) Overlap of statistically significant (Bonferroni-corrected) age-associated sites among different *in vivo* (human blood) and *in vitro* (cultured fibroblasts) studies. Note that there are more significant CpGs in cultured cells systems compared to human studies, likely due to the controlled experimental conditions *in vitro*. Only 35 CpGs overlap between the three datasets. (**B-D**) Proportions of the 35 replicated age-associated CpGs that undergo hyper- or hypo-methylation, (**B**) their position relative to CpG islands, and (**C**) and gene regions. See Supplemental Table S1 for complete annotation. (**E**) Of the 35 overlapping CpGs, 7 mapped to a gene that was differentially expressed (>0.2 fold change in transcript levels) measured by RNAseq. (**F**) Color coded DNAm levels for the CpG that map to each gene shown in (E).

### Supplemental Figure S5

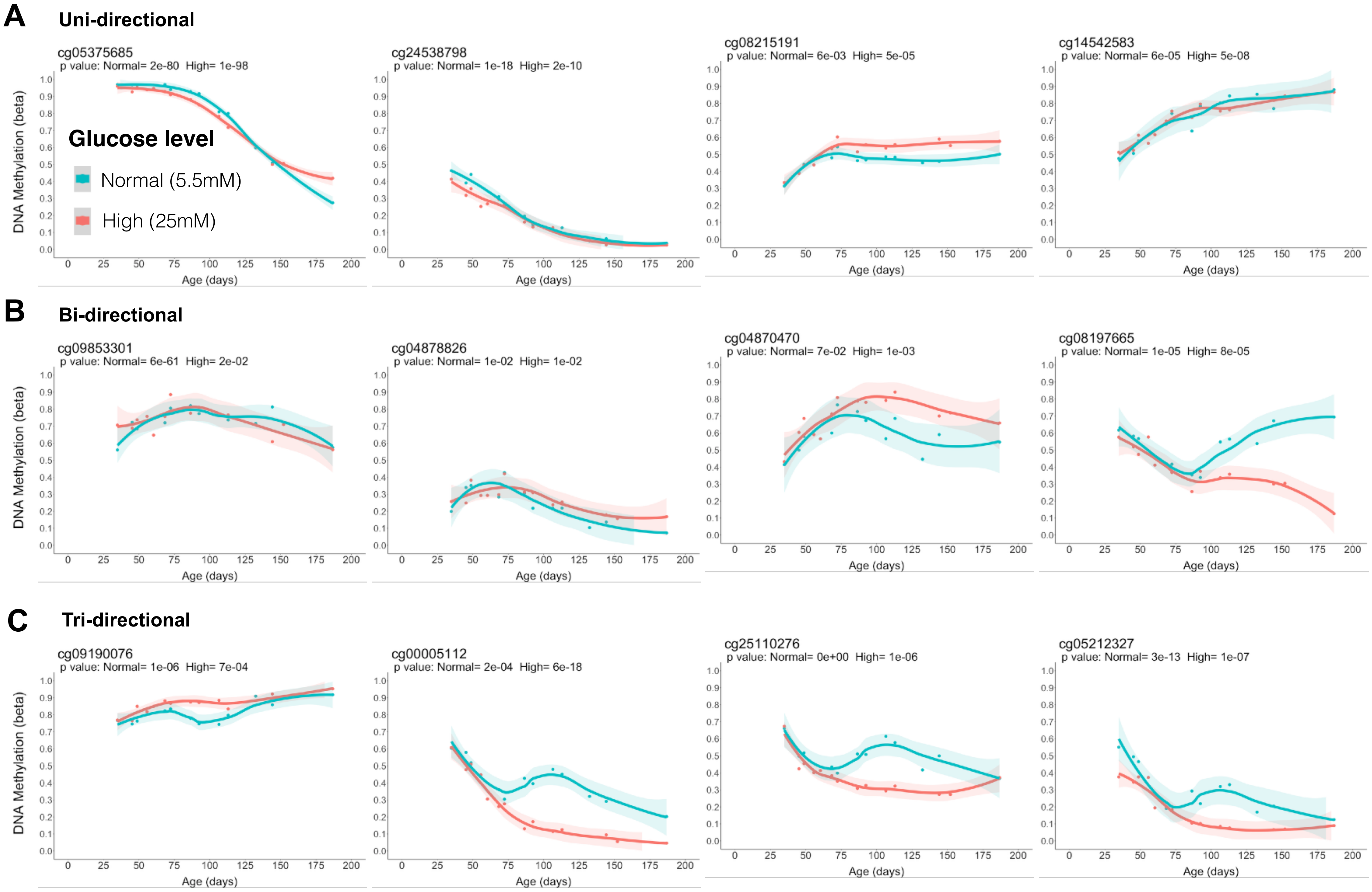


**S5. Exploratory analysis of non-monotone CpG trajectories.**

(**A**) DNAm kinetics across the cellular lifespan for CpG sites exhibiting monotone, (**B**) bi-tone, and (**C**) tri-tone trajectories. Each CpG is shown for cells aging under either normal and high glucose. Non-monotone sites were handpicked form a list of GAM-fitted sites with up to 9 DoF.

### Supplemental Figure S6 - Part 1

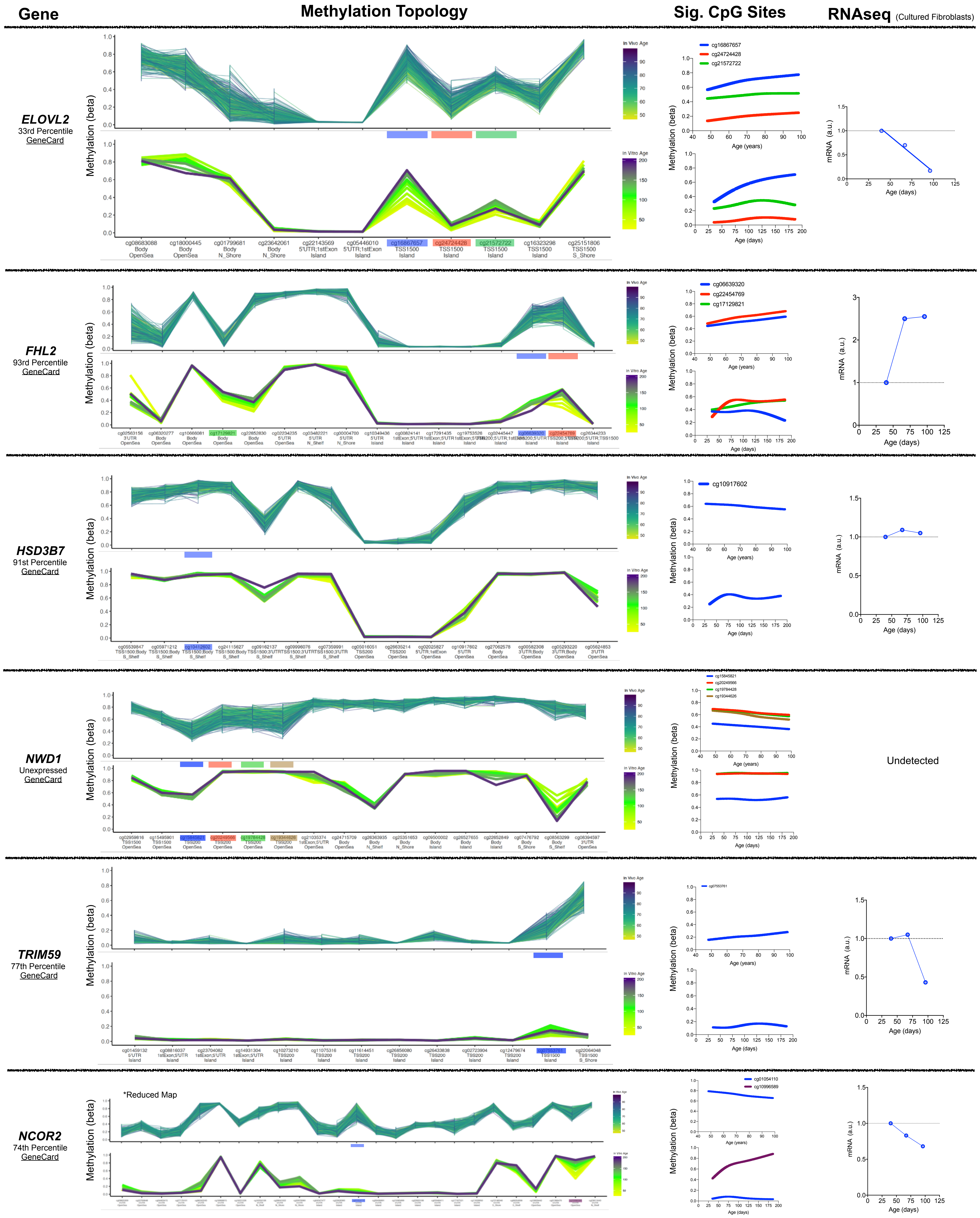

### Supplemental Figure S6 - Part 2

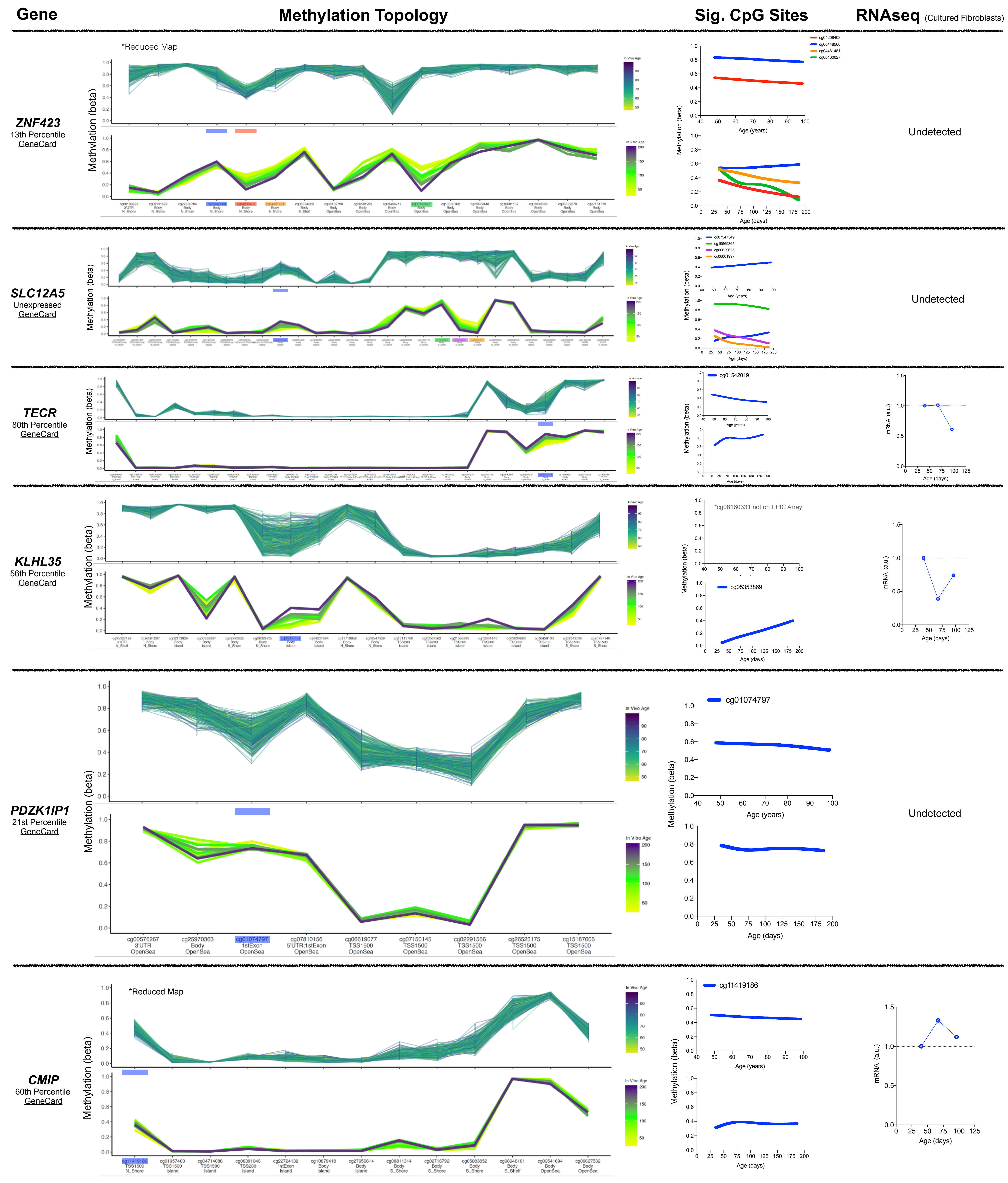

### Supplemental Figure S6 - Part 3

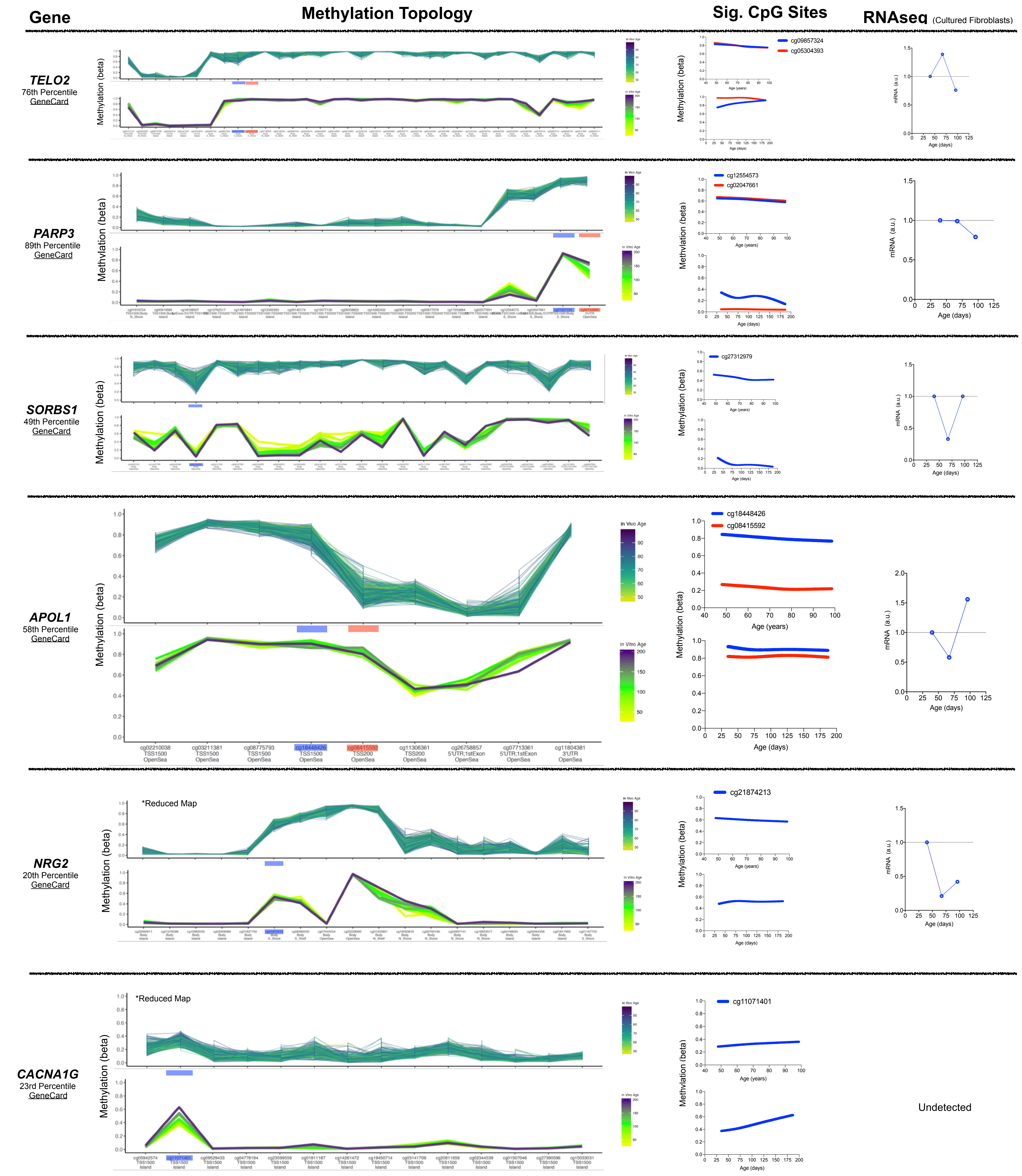

### Supplemental Figure S6 - Part 4

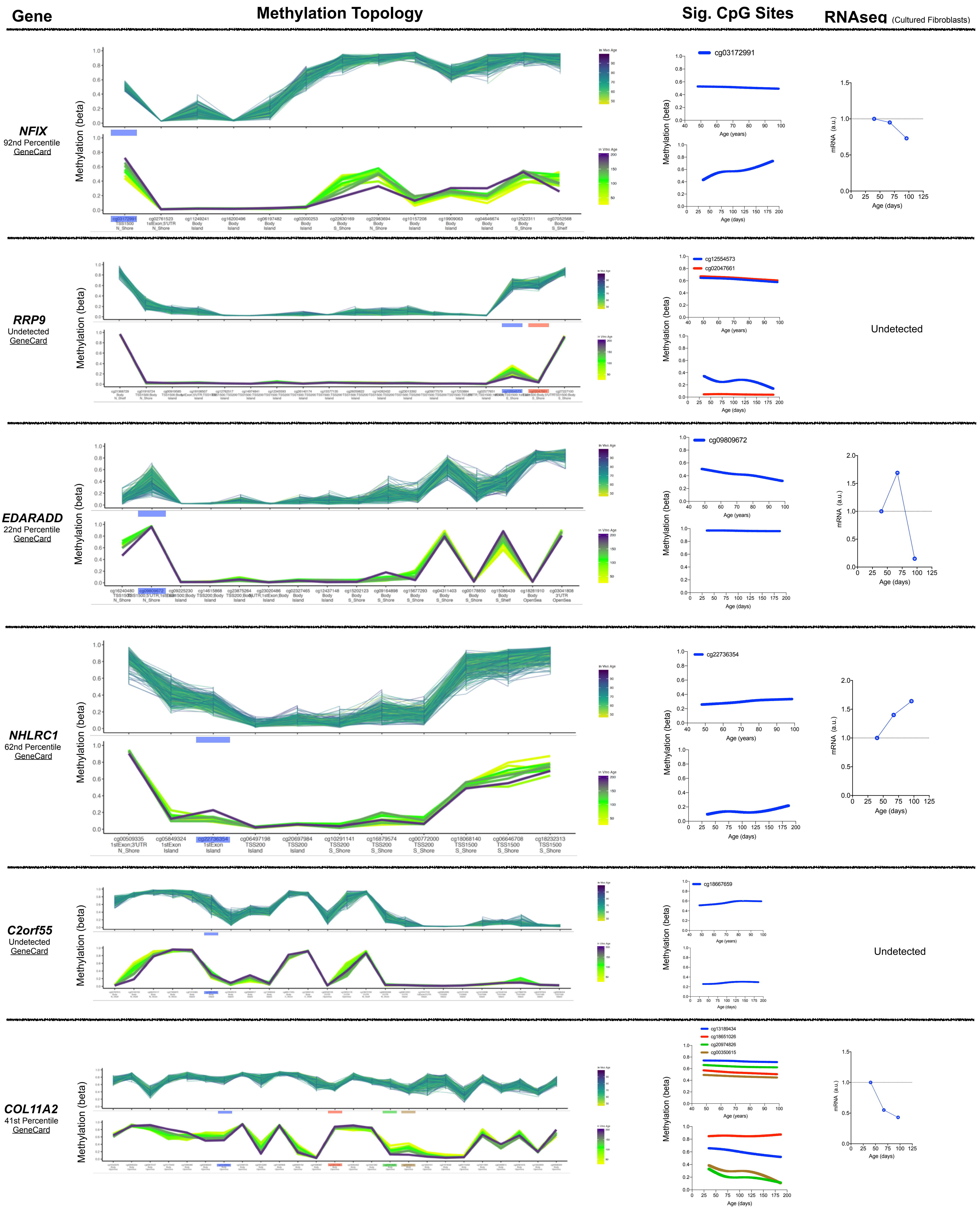

### Supplemental Figure S6 - Part 5

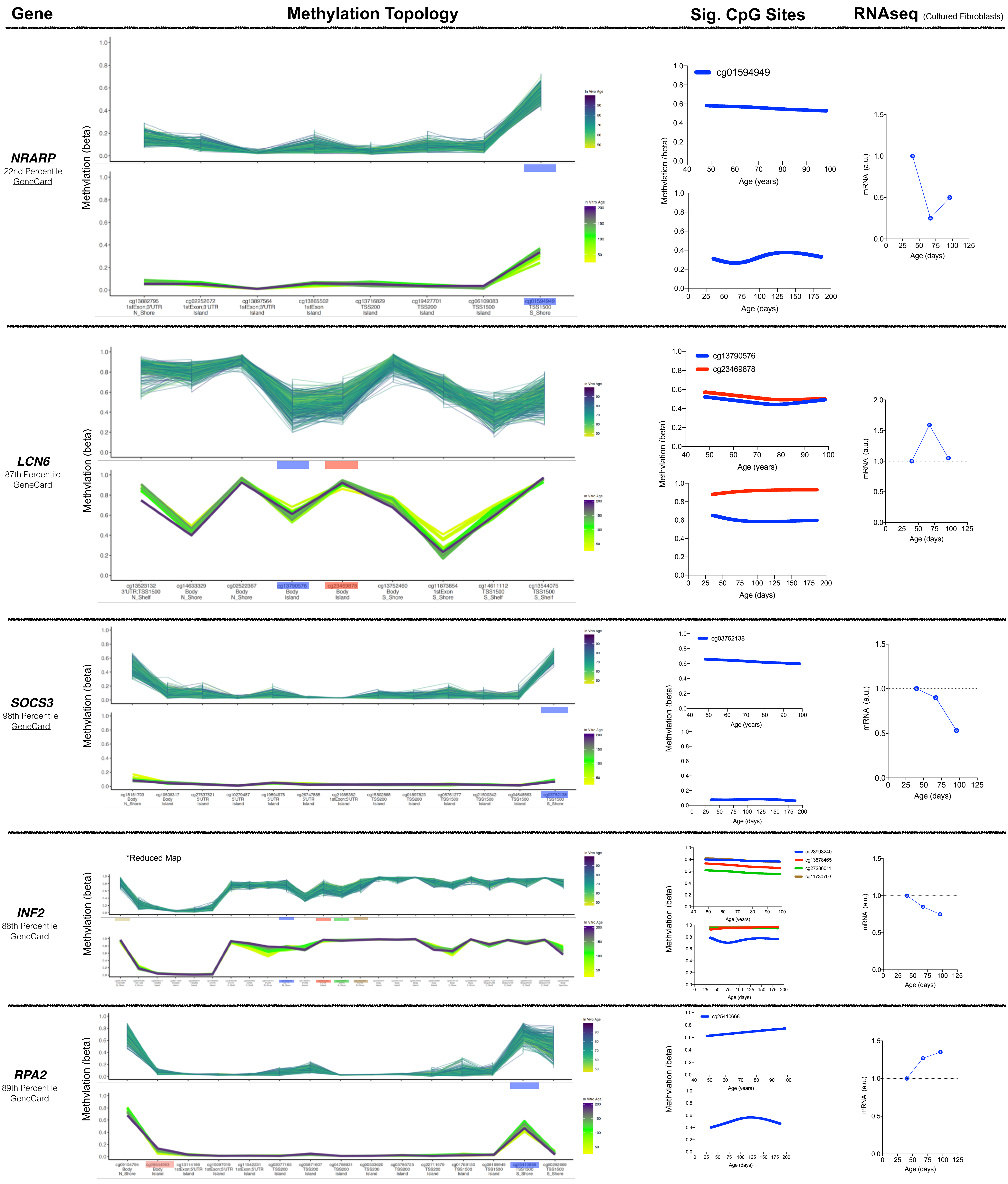


**Figure S6. Age-related DNAm gene topology graphs illustrate the degree of conservation between human blood and cultured fibroblasts.**

Detailed CpG-by-CpG gene topology maps for all 29 genes reporting at least one significant age-related CpG in the 20-year human blood longitudinal study by Wang et al. (2018). The first gene shown is *ELOVL2* for reference, as in Figure 2. The percentile shown indicates the average transcript level for each gene relative to all other transcript: 1st percentile indicates high expression, 99th percentile indicates low expression.
